# Genomic characterization and molecular evolution of human Monkeypox viruses

**DOI:** 10.1101/2022.10.17.512603

**Authors:** Patrícia Aline Gröhs Ferrareze, Rute Alves Pereira e Costa, Claudia Elizabeth Thompson

**Affiliations:** Graduate Program in Health Sciences, Universidade Federal de Ciências da Saúde de Porto Alegre (UFCSPA), Porto Alegre, RS, Brazil; Sociedade Brasileira de Valorização das Sociedades Médicas (SOBEMED), São Paulo, Brazil; Department of Pharmacosciences, Universidade Federal de Ciências da Saúde de Porto Alegre (UFCSPA), Porto Alegre, RS, Brazil

**Keywords:** Monkeypox virus, molecular evolution, genomic synteny, biological process, immunomodulatory functions, host-pathogen interaction

## Abstract

Monkeypox virus is a member of the Poxviridae family, as variola and vaccinia viruses, presenting a linear double-strand DNA genome approximately ≈197 kb long, which encodes ≈190 non overlapping ORFs. Genomic comparison of Central and West African clades shows the presence of unique genes that promote different disease presentations according to the strain. Since the last smallpox vaccination efforts ended in the mid-1980s, nowadays, there is concern about the recent spread of human monkeypox disease around the world. Currently, almost 70,000 human monkeypox cases are diagnosed in the world, of which more than 7,800 are from Brazil. This study aims to evaluate genomic epidemiology and molecular evolution of hMpxV genomes. Using computational biology to analyze 604 hMpxV genomes from 1960 up to 2022, it was possible to observe synteny breaks and gene conservation between Central and West clade genomes, with the presence of strains associated with the 2022 outbreak assigned to West African clade. Evidence of diversifying selective pressure on specific sites from protein coding sequences acting on immunomodulatory processes was identified. The existence of different sites under diversifying - and purifying - selection in paralog genes denotes adaptation mechanisms underlying the host-pathogen interaction of Monkeypox virus in human species.

**HIGHLIGHTS:**

- Synteny breaks were identified among West and Central African genomes with sequence identity of 96.5%
- Positive selection evidence was found on sites from genes of immunomodulatory functions
- Different sites under diversifying and purifying selection were observed in paralog genes
- Genomes from the 2022 outbreak are phylogenetically assigned to the West African clade

## 1 INTRODUCTION

Human Monkeypox virus (hMpxV) is a member of the family Poxviridae, genus Orthopoxvirus, as variola virus (smallpox), vaccinia virus, mousepox virus, and cowpox virus. With a massive vaccination strategy, smallpox was declared as eradicated by OMS in 1980, leaving no animal or environmental viral reservoirs (Weaver & Isaacs, 2008). In this way, last smallpox vaccination efforts ended in the mid-1980s, bringing, nowadays, concern about the recent spread of human monkeypox disease around the world. Since the first reported human monkeypox case in a child from the Democratic Republic of Congo, in 1970, some sparse occurrences have been reported in the world, such as the 2003 US smallpox outbreak (Hammarlund et al., 2005).

The hMpxVs genome is a linear double-strand DNA genome approximately ≈197 kb long, which encodes ≈190 non overlapping ORFs, besides covalently closed hairpin ends with inverted terminal repeats (ITR) at each end. The highly conserved central region encodes genes involved in transcription, replication, and virion assembly. The genes encoded in the terminal regions can vary among poxviruses, coding proteins involved in host range determination and pathogenesis (Kugelman et al., 2014). The genomic comparison of strains from Central Africa (ZAI-96) and West Africa (SL-V70, COP-58, and WRAIR-61) shows the presence of 170 shared orthologs among the strains, with a protein level identity of 99.4%, besides of the many unique genes from each group (Weaver & Isaacs, 2008). With remarkable clinical differences, West African strains promote a milder disease presentation with lower fatality and lower human-to-human transmission. The analysis of virulence genes showed the presence of 53 shared genes among strains, of which BR-203 (Virulence protein), BR-209 (IL-1b-binding protein), and COP-C3L (Inhibitor of complement enzymes) presented the most significant differences (Weaver & Isaacs, 2008).

Currently, almost 70,000 human monkeypox cases are diagnosed in the world, of which more than 7,800 are from Brazil (https://ourworldindata.org/monkeypox, accessed on October 06, 2022). With an increasing number of cases and spread patterns not completely elucidated, the genomic epidemiology and molecular evolution analyses are powerful tools to identify evolutionary changes that can potentially affect the host-pathogen interaction. Thereby, this study aims to identify the phylogenetic patterns related to the main immunomodulatory proteins coded by monkeypox virus genomes and their sites under diversifying selection.

## 2 MATERIAL AND METHODS

### 2.1 hMpxV genomes retrieval, ORF prediction and orthogroups clustering

A number of 604 complete human Monkeypox virus genomes were retrieved from the GISAID database (https://www.epicov.org/epi3/) on July 26, 2022. The ORF prediction for each genome sequence was performed with Linux x64 local version of ORFfinder (https://www.ncbi.nlm.nih.gov/orffinder/) and default parameters.

Ortholog clusterization was performed with OrthoFinder v.2.5.4 (Emms & Kelly, 2019) and default parameters. Coding sequences (CDS) from human Monkeypox virus strain Zaire_1979-005 (GenBank accession DQ011155.1 / GISAID EPI_ISL_13053218) reference genome (Likos et al., 2005) and NC_063383.1 from human Monkeypox virus strain M5312_HM12_Rivers were added in order to identify orthogroups related to known Monkeypox virus annotated genes from Central and West African clades, respectively. For multicopy orthogroups, the longest predicted ORF from each genome was selected. Multiple sequence alignments (msa) were performed with Nextalign v0.2.0 (Aksamentov et al., 2021) using the Monkeypox virus strain Zaire_1979-005 CDS sequences belonging to the Central clade as reference. The nucleotide insertions were removed from msa files.

Functional annotation was implemented with BLAST2GO (Gotz et al., 2008) and REVIGO (Supek et al., 2011) with a medium result list (0.7), removal of obsolete GO terms and SimRel semantic similarity measure. CDS visualization and genome sequence alignment for synteny evaluation were performed with Geneious Prime software (https://www.geneious.com/) using Mauve progressive aligner plugin (Darling et al., 2004) with DQ011155.1 and NC_063383.1 reference genomes.

### 2.2 Phylogenetic analyses and molecular evolution tests

For phylogenetic analyses, the previously performed msa for 155 MpxV CDS genes were selected. For each data set, the evolutionary models and maximum likelihood phylogenetic trees were inferred by IQTREE v1.6.12 (Nguyen et al., 2014; Kalyaanamoorthy et al., 2017) with an approximate Bayes test and a SH-like approximate likelihood ratio test with 1,000 replicates.

Pervasive adaptive and purifying selection tests were performed with FUBAR (Fast, Unconstrained Bayesian AppRoximation for Inferring Selection) (Murrell et al., 2013), and FEL (Fixed Effects Likelihood) (Kosakovsky Pond & Frost, 2005) methods from the HyPhy package.

### 2.3 Phylogenomic analysis

For phylogenomic analysis, the 155 CDS msa files previously obtained were selected and concatenated in a unique multiple sequence alignment file with *catfasta2phyml* Perl script (https://github.com/nylander/catfasta2phyml). Since not all sequences had orthologs identified in all hMpxV genomes, gaps were introduced when a gene sequence was absent. Additionally, the CDS alignment was previously performed against Monkeypox virus strain Zaire_1979-005 reference CDS, excluding nucleotide insertions from the sequence alignments. The evolutionary models and maximum likelihood phylogenetic trees were inferred by IQTREE v1.6.12 (Nguyen et al., 2014; Kalyaanamoorthy et al., 2017) with an approximate Bayes test and a SH-like approximate likelihood ratio test with 1,000 replicates in a non-partitioned sequence set.

Finally, a phylogenomic tree was inferred in a partitioned analysis with the coordinates of the 155 coding sequences in the multiple sequence alignment file. The evolutionary model was inferred for each partition (*-spp* parameter for partition-specific rates) and the tree was rebuilt using IQTREE v1.6.12 (Nguyen et al., 2014; Kalyaanamoorthy et al., 2017) with an approximate Bayes test and a SH-like approximate likelihood ratio test with 1,000 replicates. Temporal sign of the samples was evaluated with TempEst v.1.5.3 (Rambaut et al., 2016).

## 3 RESULTS

### 3.1 Genomic alignment and synteny evaluation

The comparison of Central African clade genomic sequence DQ011155.1 (human Monkeypox virus strain Zaire_1979-005) and West African clade genomic sequence NC_063383.1 (human Monkeypox virus strain M5312_HM12_Rivers) indicates the occurrence of two major synteny breaks in these genomes. The first one, in the Central African Monkeypox virus strain Zaire_1979-005 contains the coding sequences for genes MPXV_ZAI1979_005_022, MPXV_ZAI1979_005_023, MPXV_ZAI1979_005_024 and MPXV_ZAI1979_005_025 that code for complement control protein AAY97217, and kelch-like proteins AAY97218, AAY97219 and AAY97220, respectively. The second one, located on West African Monkeypox virus strain M5312_HM12_Rivers do not contain any annotated gene.

Moreover, other small synteny breaks between the collinear blocks from both genomes can be identified. Genes MPXV_ZAI1979_005_008 (unknown protein AAY97203), 009 (unknown protein AAY97204), 010 (unknown protein AAY97205), 012 (interleukin-1 receptor antagonist-like protein AAY97207), 057 (unknown protein AAY97252), 058 (unknown protein AAY97253), 142 (cowpox A-type inclusion protein AAY97337), 143 (cowpox A-type inclusion protein AAY97338), 144 (cowpox A-type inclusion protein AAY97339), 145 (cowpox A-type inclusion protein AAY97340), 167 (unknown protein AAY97362), 168 (unknown protein AAY97363), 174 (kelch-like protein AAY97367), 189 (bifunctional IL-1-beta-inhibitor AAY97381), 189.5 (bifunctional IL-1-beta-inhibitor AAY97382), 190 (unknown protein AAY97383), 191 (unknown protein AAY97384), 194 (kelch-like protein AAY97387), 195 (serine protease inhibitor-like SPI-1 protein AAY97388), 198 (TNF-alpha-receptor-like protein AAY97392), 199 (unknown protein AAY97393), and 201 (unknown protein AAY97395) have no orthologues in NC_063383.1 strain according to genome collinearity analysis. The same is observed for genes OPG059 (cytochrome C oxidase), OPG076 (MV membrane EFC component), OPG158 (Viral membrane assembly proteins A32.5L) in comparison with the DQ011155.1 genome (Supplementary File 1). The pairwise genomic sequence identity is calculated to 96.5%.

### 3.2 ORF prediction and orthogroup assignment

Whole genome ORF prediction resulted in 1,027,404 annotated ORFs from 604 hMpxV genomes, with an average mean of 1,701 predicted ORFs by genome. Of these, 501,999 ORFs were predicted in the complementary strand. The ortholog groups assignment clustered 1,024,871 ORFs (99.7%) in 1,617 orthogroups. There were 103 orthogroups with all species present (n = 604) and 6 of these consisted entirely of single-copy genes. In each orthogroup, the longest ORF was selected as representative for each genome.

Using Central African Monkeypox virus strain Zaire_1979-005, which encodes 202 sequences (n = 202), as reference for comparison among all 604 genomes, 200 orthogroup sequence sets were identified. Subsequently, 151 were selected since their sequences could be aligned to both Central African Monkeypox virus strain Zaire_1979-005 reference coding sequences (n = 202) and West African Monkeypox virus strain M5312_HM12_Rivers (n = 179) reference coding sequences. Moreover, these 151 orthogroups had predicted orthologous ORFs identified in ≥302 genomes (50%). For151 orthogroups, 155 genes and their respective ortholog sequences were retrieved (Figure 2). A total of 42.58% of these 155 genes had ortholog sequences identified in the 604 hMpxV genomes. Thirty-nine orthogroups presented nucleotide insertions in ≥10 sequences in comparison with Central African Monkeypox virus strain Zaire_1979-005 reference coding sequences; 534 hMpxV samples presented nucleotide insertions in their genes from all 39 orthogroups, generating possible frame shifts in these gene ORFs. Stop codons and nucleotide insertions were deleted or disregarded in the sequence alignment step. Only positions aligned to the reference CDS were evaluated by molecular evolution tests and phylogenomic analyses.

**Figure 1.**
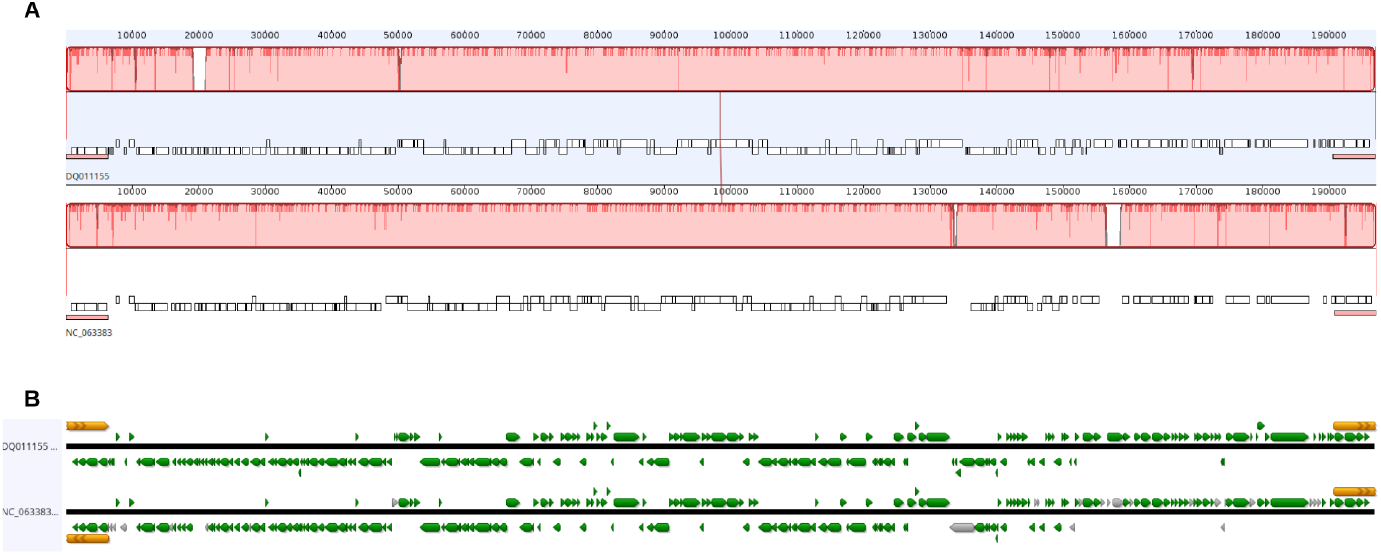
Genomic alignment and synteny evaluation between Central African clade sequence DQ011155.1 (human Monkeypox virus strain Zaire_1979-005) and West African clade sequence NC_063383.1 (human Monkeypox virus strain M5312_HM12_Rivers). (A) Synteny breaks (white spaces) between collinear genomic blocks (blocks). (B) Locus correspondence between aligned genomes in both DNA strands. Annotated genes are highlighted in green. Terminal repeat regions are highlighted in orange.

**Figure 2.**
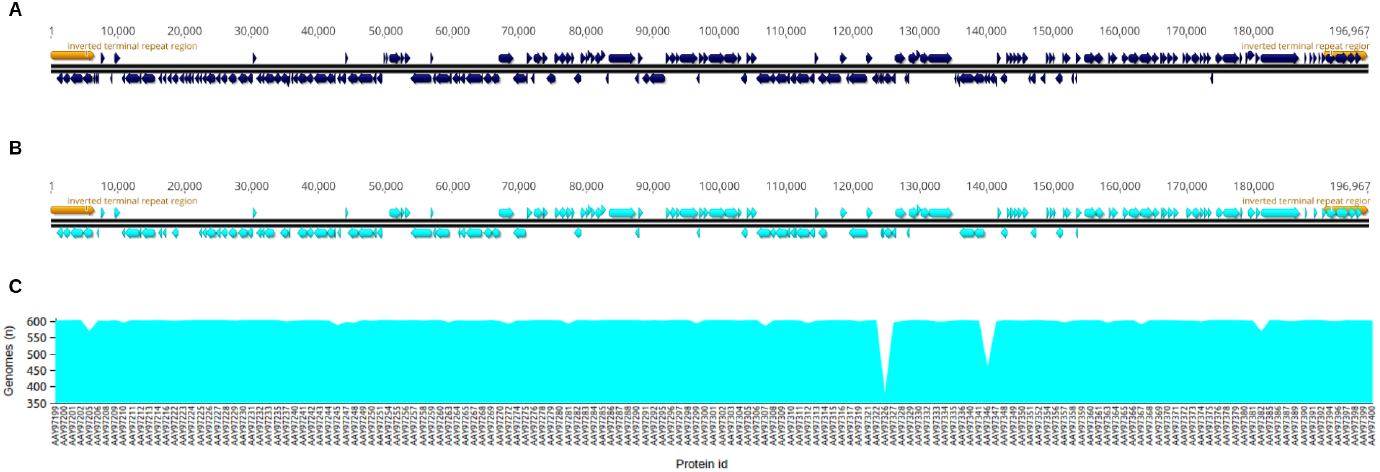
Coding sequences from genes annotated for Monkeypox virus strain Zaire_1979-005. (A) The 202 annotated coding sequences from Monkeypox virus strain Zaire_1979-005. (B) The 155 coding sequences related to 151 selected orthogroups identified in this study. (C) Number of genomes with identified orthologs for the 155 selected coding sequences; the “y” axis indicates the number of sequences and the “x” axis indicates the protein ID related to the identified CDS.

Despite 151 orthogroups were annotated by OrthoFinder, four of them included two paralogous genes each. These two orthogroups belong to coding sequences from chemokine-binding proteins AAY97199 (complement: 884 - 1624) / AAY97400 (195344 - 196084); TNF-alpha-receptor-like proteins AAY97200 (2,798 - 1,752) / AAY97399 (194,170 - 195,216); and ankyrin-like proteins AAY97201 (4,651 - 2,888) / AAY97398 (192,317 - 194,080) and AAY97202 (complement: 4,808 - 6,121) / AAY97397 (190,847 - 192,160). In this way, the differentiation of the true orthologs relative to each CDS was performed according to the CDS and predicted ORF location in genomes, in order to maintain genomic synteny. No nucleotide mutations or size alterations were observed between these paralogs. Both of them are located in the opposite Inverted Terminal Repeat Regions (ITRs) in order that each ITR presents a copy of each gene.

The analysis of the Gene Ontology terms related to the 155 genes included in this study (from 151 orthogroups) showed the prevalence of viral processes (Figure 3), including modulation by virus of host process and viral life cycle.

**Figure 3.**
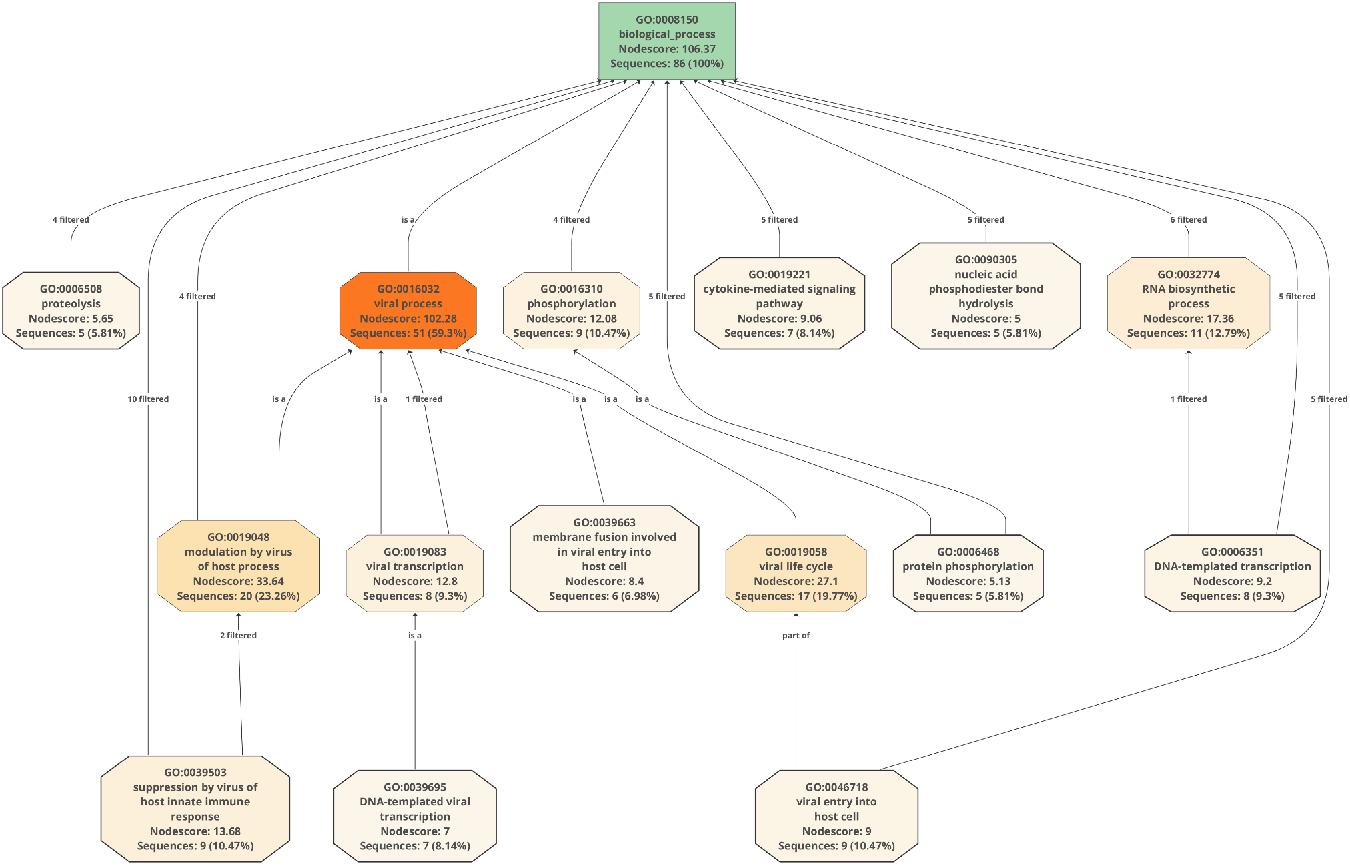
Main biological processes associated with the 151 annotated orthogroups. Search performed with Monkeypox virus strain Zaire_1979-005 reference coding sequences used in this study.

### 3.3 Molecular Evolution

The application of the Hyphy method FUBAR identified positive adaptive selection evidence in 33 genes (Figure 4 and Table 1), of which 5 were associated to biological processes of immunomodulatory functions of suppression of immune response by inhibition of IRF3 activity and type I interferon-mediated signaling pathway, as well suppression of antigen processing and presentation of peptide antigen via MHC class II (Figure 5 and Table 1). Other three genes are related to viral entry in host cells, viral material transportation, and host protein ubiquitination (Table 1). Among all gene ontology terms related to these genes, *Suppression by virus of host innate immune response* (GO:0039503) and *Viral DNA genome replication* (GO:0039693) were the most frequently annotated biological processes despite not the most enriched terms (Figure 4).

**Table 1.** Positively selected sites identified by FUBAR method in hMpxV protein coding sequences.

| DQ011155.1 |  | NC_063383.1 |  | Biological Process | Domain?<br>(Superfamily) | Site | FUBAR |  |  |
| --- | --- | --- | --- | --- | --- | --- | --- | --- | --- |
| Locus<br>MPXV_<br>ZAI1979<br>_005 | Protein<br>AAY | Locus<br>NBT03_<br>gp | Gene<br>OPG |  |  |  | Alpha | Beta | Post<br>prob |
| _004 | 97202 | 004 | 015 | -- | PHA02795 | 312 | 4.482 | 36.464 | 0.9333 |
| _010 | 97205 | -- | -- | Viral process | PHA02725 | 62 | 3.947 | 24.712 | 0.9057 |
|  |  |  |  |  | PHA02725 | 64 | 3.913 | 28.960 | 0.9242 |
| _015 | 97210 | 008 | 023 | -- | PHA02917 | 258 | 3.738 | 34.489 | 0.9423 |
|  |  |  |  |  | PHA02917 | 426 | 3.373 | 34.249 | 0.9485 |
| _017 | 97212 | 010 | 025 | -- | PHA02792 | 255 | 3.637 | 24.691 | 0.9140 |
| _018 | 97213 | 011 | 027 | Suppression by virus of host innate immune response | PHA02919 | 85 | 2.536 | 30.287 | 0.9556 |
| _028 | 97223 | 017 | 036 | Suppression by virus of host viral-induced cytoplasmic pattern recognition receptor signaling pathway via inhibition of IRF3 activity | Poxvirus | 25 | 3.581 | 30.392 | 0.9339 |
| _029 | 97224 | 018 | 037 | -- | PHA02946 | 16 | 4.350 | 36.237 | 0.9352 |
| _031 | 97226 | 020 | 039 | Suppression by virus of host innate immune response | PHA02791 | 256 | 4.190 | 30.127 | 0.9192 |
| _038 | 97233 | 027 | 047 | -- | PHA02790 | 473 | 2.184 | 31.477 | 0.9627 |
|  |  |  |  |  | PHA02790 | 476 | 3.706 | 25.479 | 0.9123 |
|  |  |  |  |  | PHA02790 | 478 | 4.439 | 34.560 | 0.9313 |
| _046 | 97241 | 035 | 055 | -- | PHA02699 | 319 | 3.698 | 31.946 | 0.9362 |
| _054 | 97249 | 044 | 064 | Microtubule-dependent intracellular transport of viral material towards cell periphery | PHA02695 | 88 | 3.666 | 32.970 | 0.9385 |
|  |  |  |  |  | PHA02695 | 485 | 4.465 | 33.039 | 0.9228 |
| _060 | 97255 | 048 | 069 | -- | E7R | 38 | 3.510 | 30.287 | 0.9331 |
| _062 | 97257 | 050 | 071 | DNA replication; DNA recombination; Viral DNA genome replication; DNA biosynthetic process; Nucleic acid phosphodiester bond hydrolysis | PHA03036 | 785 | 4.202 | 35.889 | 0.9368 |
| _065 | 97260 | 053 | 074 | -- | PHA02693 | 515 | 2.498 | 35.993 | 0.9687 |
| _074 | 97269 | 063 | 084 | -- | PHA02653 | 428 | 2.375 | 33.498 | 0.9650 |
|  |  |  |  |  | PHA02653 | 432 | 3.128 | 32.911 | 0.9487 |
|  |  |  |  |  | PHA02653 | 441 | 3.769 | 35.854 | 0.9442 |
| _083 | 97278 | 072 | 093 | Regulation of DNA-templated transcription; DNA-templated viral transcription | Pox_TAP | 30 | 3.255 | 35.211 | 0.9528 |
| _084 | 97279 | 073 | 094 | Fusion of virus membrane with host plasma membrane; Viral entry into host cell | Pox_G9-A16 | 194 | 3.612 | 33.409 | 0.9415 |
| _095 | 97290 | 084 | 105 | DNA-templated transcription | RNAP_largest_subunit_N | 337 | 3.292 | 34.932 | 0.9505 |
| _103 | 97298 | 092 | 113 | Polynucleotide 5' dephosphorylation; RNA 5'-cap (guanine-N7)-methylation; 7-methylguanosine mRNA capping | NO | 3 | 3.966 | 35.897 | 0.9409 |
|  |  |  |  |  | NO | 543 | 3.964 | 35.584 | 0.9401 |
| _105 | 97300 | 094 | 115 | -- | Pox_D3 | 171 | 2.535 | 25.878 | 0.9007 |
| _107 | 97302 | 096 | 117 | DNA replication, synthesis of RNA primer; Viral DNA genome replication | NO | 685 | 4.455 | 33.757 | 0.9246 |
| _108 | 97303 | 097 | 118 | -- | NO | 4 | 3.762 | 31.903 | 0.9349 |
| _120 | 97315 | 109 | 130 | -- | Orthopox_A5L | 63 | 2.876 | 35.851 | 0.9611 |
| _155 | 97350 | 141 | 163 | Suppression by virus of host antigen processing and presentation of peptide antigen via MHC class II | Chordopox_A35R | 46 | 4.060 | 34.490 | 0.9354 |
| _162 | 97357 | 148 | 172 | -- | Orthopox_A43R | 63 | 4.267 | 31.832 | 0.9241 |
| _166 | 97361 | 152 | 176 | -- | NO | 4 | 4.374 | 35.391 | 0.9319 |
| _174 | 97367 | -- | -- | Modulation by virus of host protein ubiquitination;<br>Modulation by virus of host ubiquitin-protein ligase activity | NO | 2 | 3.619 | 21.985 | 0.9137 |
|  |  |  |  |  | NO | 3 | 3.664 | 28.575 | 0.9314 |
|  |  |  |  |  | NO | 4 | 3.742 | 22.907 | 0.9060 |
| _186 | 97378 | 166 | 198 | Protein phosphorylation | PKc_like | 87 | 1.760 | 24.985 | 0.9124 |
| _192 | 97385 | 169 | 204 | Suppression by virus of host type I interferon-mediated signaling pathway | NO | 235 | 6.032 | 37.459 | 0.9007 |
| _193 | 97386 | 170 | 205 | -- | PHA03095 | 689 | 3.731 | 38.212 | 0.9545 |
|  |  |  |  |  | PHA03095 | 713 | 6.977 | 41.756 | 0.9018 |
| _197 | 97391 | 173 | 210 | -- | NO | 1583 | 4.172 | 33.519 | 0.9306 |
| _203 | 97397 | 176 | 015 | -- | PHA02795 | 242 | 4.618 | 35.530 | 0.9283 |
| _204 | 97398 | 177 | 003 | -- | PHA03095 | 68 | 3.366 | 30.274 | 0.9388 |
|  |  |  |  |  | PHA03095 | 263 | 3.439 | 33.356 | 0.9461 |
|  |  |  |  |  | PHA03095 | 567 | 3.890 | 35.275 | 0.9428 |

Domain? indicates if the positively selected site is located on a protein domain or outside ("NO"). Gray rows indicate sites identified by the FEL method. Red lines indicate potentially false positive results

**Figure 4.**
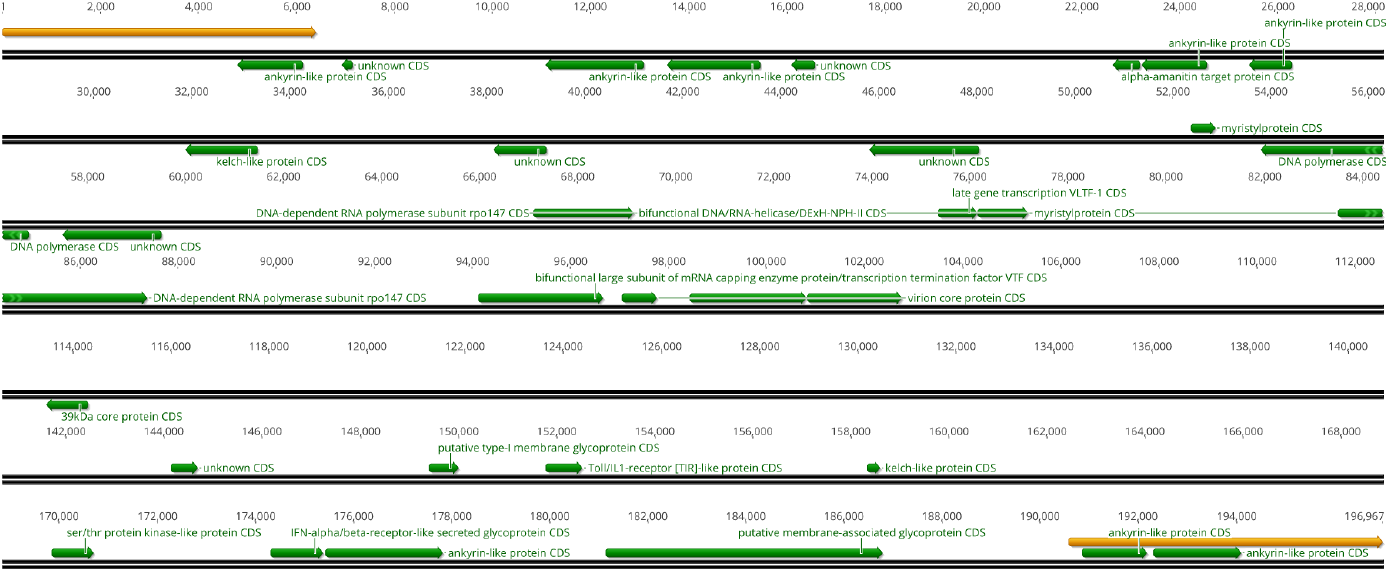
Genomic visualization of the CDS associated with the positively selected sites identified by HyPhy methods.

**Figure 5.**
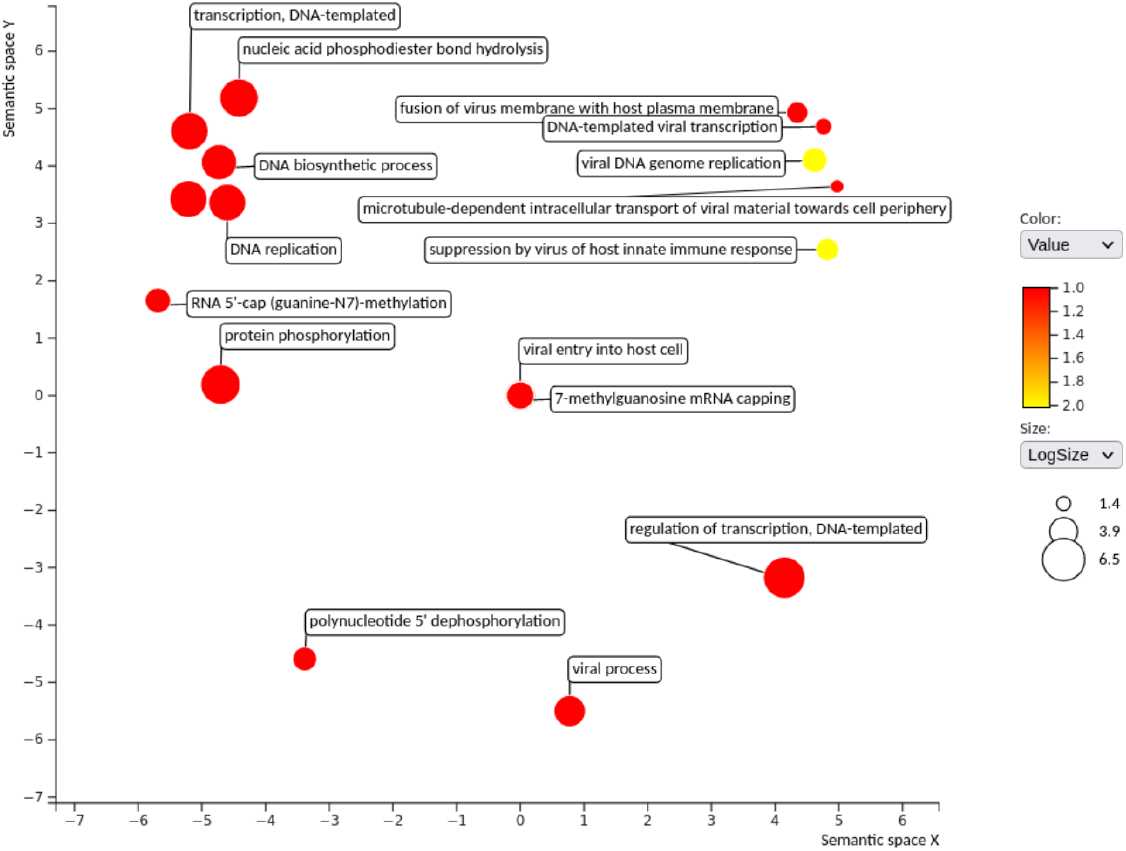
Biological process enrichment analysis from sequences with sites under adaptive selection. Color gradient indicates GO terms frequency in these gene annotations. Log size indicates the GO terms enrichment.

No sites under adaptive selection were inferred by FEL besides those previously identified by FUBAR method. Interestingly, of the 33 sequences, 7 are described as ankyrin-like proteins. The Monkeypox virus strain Zaire_1979-005 codifies 11 ankyrin-like proteins, of which 8 present evidence of pervasive diversifying selection in our analysis and two of them are from paralogous genes (AAY97202 / AAY97397). These two paralog genes have different sites with adaptive selection evidence (312 and 242). Both sites are in the PHA02795 superfamily domain and resulted from a nucleotide substitution in both first and second codon bases, altering a polar tyrosine by a non-polar valine (AAY97202) and a negatively charged glutamic acid by a non-polar leucine (AAY97397) (Figure 6). The second akyrin-kike protein paralog copy AAY97398 presents three sites under positive selection: a cysteine to a histidine in codon 68, asparagine to aspartic acid in codon 263, and lysine to tryptophan in codon 567. Similarly, the MpxV genome codifies 7 kelch-like proteins, of which 2 may be under adaptive selection. Seven genes present positively selected sites outside their protein domains. Despite most of the described genes (Table 1) have no annotated biological process, it is possible to infer the molecular function by domain recognition.

**Figure 6.**
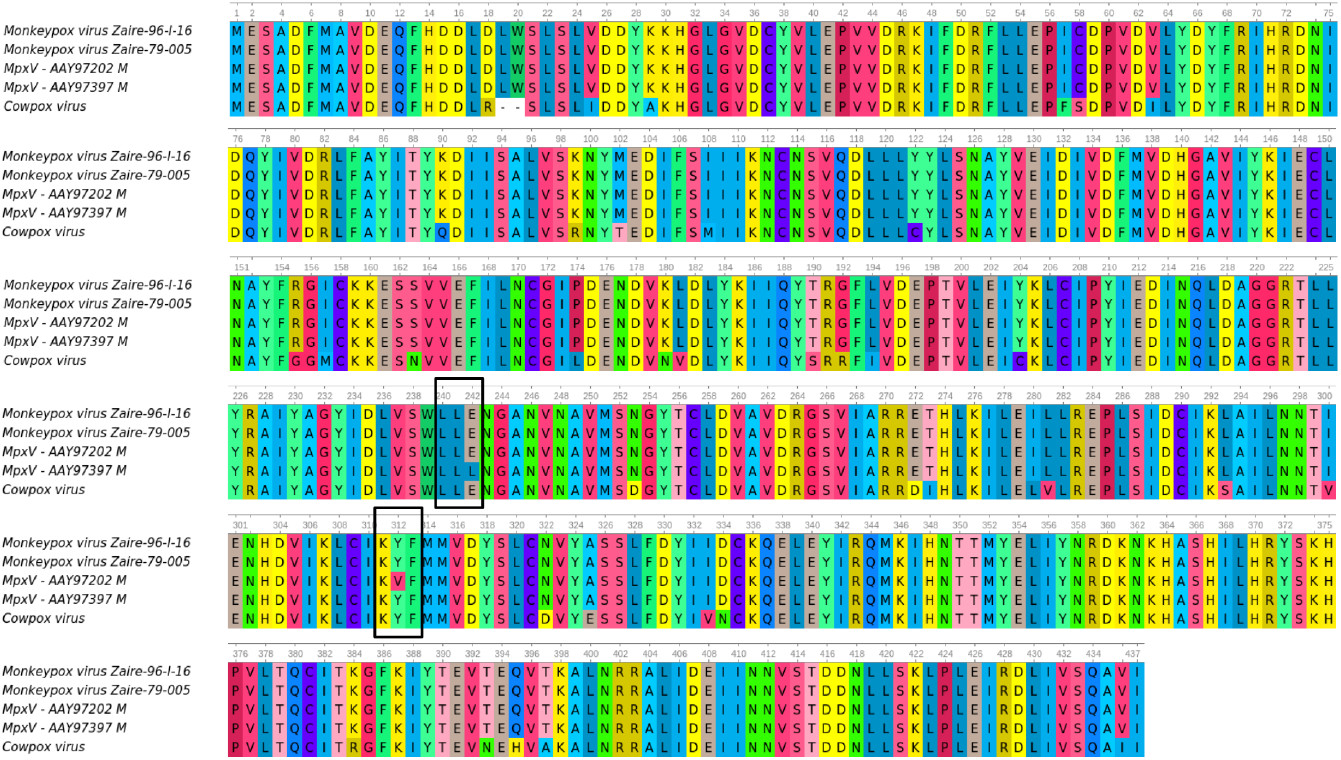
Sequence comparison of Ankyrin-like protein domain PHA02795 superfamily. CDD Search identified 100% of sequence identity between AAY97202 from Monkeypox virus Zaire-79-005 and an ortholog from Monkeypox virus Zaire 96-I-16. Since AAY97202 and AAY97397 paralogs also share 100% of sequence identity, both of them (MpxV - AAY97202 / MpxV - AAY97397) are presented with their potential mutated sites under adaptive selection, which are presented inside the black boxes.

Importantly, from the 33 coding sequences with codons presenting adaptive selection evidence, 25 present the same predicted sequence length between Central African and West African genomes. Other six sequences present small differences of one up to three additional amino acids in the coded protein. Sequences AAY97202 / AAY97397 as well as the unknown protein AAY97205 had no identified orthologues in Variola (NC_001611.1) and Vaccinia (MT946551.2) genomes (Supplementary File 2). Interestingly, protein AAY97205 has no officially annotated ortholog gene in West African genome NC_063383.1, as previously seen in genomic synteny analysis. However, in this study, an ORF for a 73 aa coded protein (AAY97205 is coded as a 65 aa sequence) was predicted for this genome, suggesting a potential non-annotated ortholog gene (Supplementary File 3A). However, the genetic dissimilarity in 3’ sequence alignment (evidenced by a small synteny break in genome alignment) could result in false positive sites under adaptive selection pressure. Similarly, positive selected sites from sequence AAY97367 could be either an artifact from a mismatched alignment. In this case, specifically, the analysis of the predicted ORFs for West African genomes and the evaluation of the collinearity between the reference genomes DQ011155.1 and NC_063383.1 shows that the predicted sequence for this kelch-like protein in West African genomes codes for approximately 144 aa while Central African potential ortholog sequences code for 82 aa (75 aa with identity of 98.67%). In this way, the first 59 aa coded by this gene in West African genomes comprises a non-syntenic region (Supplementary File 3B), creating a four amino acid sequence mismatch from the first twelve nucleotides in CDS alignment (Central African DQ011155.1 as reference).

A hundred and twenty sites were found to be under purifying selection in 64 CDS alignments evaluated in FEL analysis (Supplementary File 4). These genes were most related to biological processes of DNA transcription and DNA replication, and negative regulation/ suppression of host functions (Supplementary File 4). Interestingly, the paralogs AAY97202 and AAY97397 share three (27, 116 and 431) of the four sites with evidence of purifying selection pressure, sites 342 and 264 were differently identified for AAY97202 and AAY97397, respectively. The paralog gene couples AAY97199 / AAY97400 share site 125 and AY97200 / AY97399 share sites 31 and 221 as potentially under purifying selection. Sequence AAY97398 have no identified sites under negative selection. However, its paralogue AAY97201 presented evidence of purifying selection in three sites (256, 502 and 518).

### 3.4 Phylogenomic analysis

The phylogenomic analysis (Figure 7) using the non-partitioned concatenated multiple sequence alignment of the 155 previously selected CDS of hMpxV genomes covered 76.85% of hMpxV genomes length (in comparison with the 196,967 nt of reference genome). Initially, it is possible to observe the formation of a well structured West African clade, with 100% of branch support in SH-aLRT and approximate Bayes tests and two statistically supported subclades. The first one comprises hMpxV sequences dated to 2021/2022 (from several countries) and samples collected in 2017/2018 (England, India, Israel, Nigeria, Singapore, Thailand and USA). The second one includes older sequences from the USA (1962) and Liberia (1970). Separately, sequences collected in years 1978, 1979, 1985, 2003, 2005, 2006, and 2007, mostly from Democratic Republic of the Congo (DRC), are presented as basal groups in the tree.

**Figure 7.**
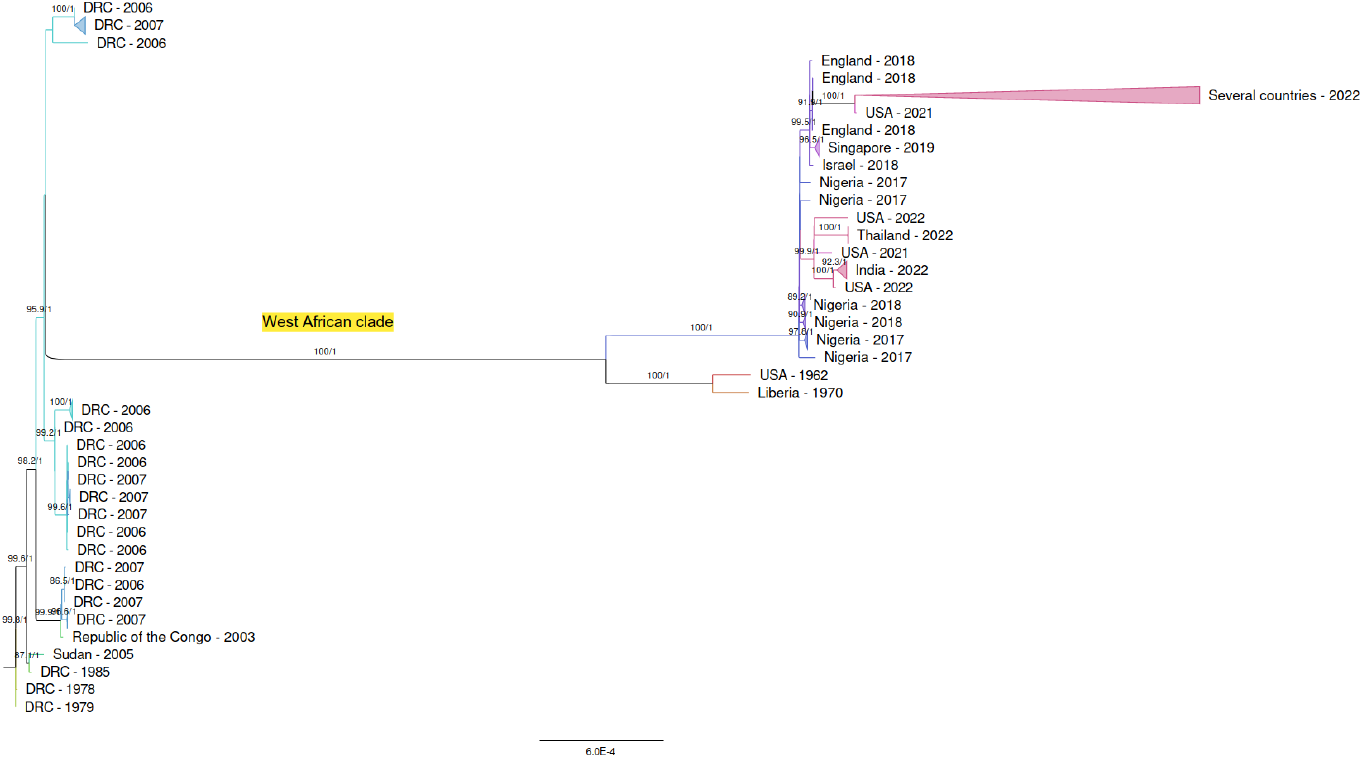
Phylogenomic analysis with 155 concatenated hMpxV gene coding sequences from 604 hMpxV genomes. Collection date and location are described in the tree. Branch support values ≥80% in both tests are labeled. Sequence groups from the same date and location were collapsed.

For the partitioned tree (Supplementary File 5A), it is possible to observe a similar phylogenetic pattern, with DRC sequences in most basal branches and a subdivision of the West African monophyletic group in two subclades. Interestingly, in both trees a hMpxV sequence from Sudan, 2005, clustered with an older DRC sequence from 1985 in a statistically supported clade (87.5/1 - 87.1/1 for SH-aLRT/aBayes test).

The analysis of the non-partitioned phylogenomic tree with TempEst software evaluated the temporal signal of the sequences. In this way, a best-fit tree was suggested (Supplementary File 5B). In this tree, Central and West African clades can be observed with different common ancestors. However, the correlation coefficient resulting from genetic divergence *versus* temporal signal was lower than the original tree (0.522 *vs*. 0.7516).

## 4 DISCUSSION

The two subfamilies of Poxviruses, *Entomopoxvirinae* and *Chordopoxvirinae*, infect a large range of species, including insects (*Entomopoxvirinae*), reptiles, birds, and mammals (*Chordopoxvirinae*). Due to this long evolutionary process of co-speciation with their hosts, host range traits are accountable for the most significant phenotypic differences between members of the *Poxviridae* family (Hendrickson et al., 2010). The orthopoxviruses present the broadest host range of the *Chordopoxvirinae*. The large coding capacity of the canonical orthopoxviral genome leads to codification of viral proteins that enable the virus to cross species barriers (Werden et al., 2008). Genome organization and synteny are consistently maintained throughout *Chordopoxvirinae* species (Hendrickson et al., 2010). Additionally, it is remarkable the conservation of many host range genes among family members (Werden et al., 2008).

Since 1970, the monkeypox disease has been observed in human hosts. Occurring in Western and Central African countries, the reported symptoms for the West African strains are milder and present lower transmission rate between humans. In fact, the case-fatality rates related to Central African strains is ∼10 % in non-vaccinated individuals (Likos et al., 2005). In 2003, the MpxV outbreak in the USA was attributed to imported rodents from Ghana,which were infected with West African strains. Despite the 37 human cases, no human-to-human transmission was reported. Conversely, the 10 monkeypox cases in the Republic of Congo showed sustained human-to-human transmission for Central African strains infection (Likos et al., 2005). Phylogenetic analyses confirm the division of monkeypox virus strains in geographically separated clades (Likos et al., 2005). Samples from Gabon, Cameroon, Republic of the Congo, and DRC belong to the Central African clade (also known as Congo Basin clade), while Nigerian, Liberian, and USA samples cluster in the West African clade (Likos et al., 2005). Since the selection of well-conserved genes to generate high-quality alignments and to estimate the evolutionary history of the *Poxviridae* family was already performed and described in literature (Hendrickson et al., 2010), we decided to apply this methodology in order to generate a reliable data set for phylogenomic analysis. The evaluation of a concatenated multiple sequence alignment covering approximately 75% of the MpxV genome presents an observable temporal signal, where genome sequences from the 2021-2022 outbreak are clustered together, with samples from 2017-2018 in the basal branches of this subclade. The formation of a monophyletic group with sequences from USA (1962) and Liberia (1970) evidentiate these sequences as belonging to the West African clade.

Numerous genetic variance mechanisms take place on Orthopoxvirus genomes evolution and co-speciation with their hosts, including those related to single nucleotide polymorphisms, small nucleotide insertions and deletions, horizontal gene transfer (HGT) and fragmentation and loss of genetic material (Hendrickson et al., 2010; Kugelman et al., 2014). Previous studies with poxviruses elucidated the host adaptation mechanism and fitness gain by recombination-mediated gene duplications and adaptive point mutations (Elde et al., 2012). Homologous genes located in terminal ends of Poxviruses are majoritarily associated with immunomodulation processes, such as the inhibition of apoptosis and antiviral pathways, potentially affecting host range determination (Gubser et al., 2004). Thus, the existence of paralogous ankyrin-like proteins and chemokine-binding proteins in ITR regions raises questions about the importance and functionality of these genes in host-pathogen interaction and immune response defenses. In this way, the detection of different sites under diversifying selection pressure, in these paralogs genes, may suggest some advantage and fitness gain in the adaptation mechanisms to the host immune system response. The (Ankyrin) ANK-repeat motif - a 33 amino acid domain found as a tandem array of 2 - 7 repeats - shows up as a mark of poxviruses, since the occurrence of protein-protein interactions mediated by theses repeats are not usually found in viruses but in bacterial, archaeal, and eukaryotic proteins acting on cell–cell signaling, cytoskeleton integrity, transcription and cell–cycle regulation, inflammatory response, and development, among others (Werden et al., 2008; Mosavi et al., 2004). Biologically relevant proteins such as the family of INK4 tumor suppressors, p15, p16, p18, and p19, as well as 53BP2, and the regulator of the tumor suppressor p53, as well as Nf-κB, contain ankyrin repeats (Mosavi et al., 2004). In fact, poxvirus proteins are more related to eukaryotic proteins than bacterial, possibly due to horizontal gene transfer from eukaryotic hosts, which may explain the acquisition of sequences with specific structural scaffolds related to immunomodulatory functions (Bahar et al., 2011).

As a result of the molecular evolution analysis, alterations were observed in genes encoding suppression by virus of host innate immune response (GO:0039503) and viral DNA genome replication (GO:0039693) proteins, as well as for ankyrin-like and kelch-like proteins. This is particularly important as it reinforces the analysis of previous studies with poxviruses elucidating the host adaptation mechanism and fitness gain (by recombination-mediated gene duplications and adaptive point mutations). In fact, as already known, ankyrins has as its main and first GO annotations, the virus receptor activity followed by DNA and RNA binding. Its role in smallpox and vaccinia viruses is also well described (Shchelkunov et al., 1993). It is no different for kelch-like proteins that have the same GO annotations, in the same order than ankyrin-like proteins. In general, kelch-like proteins are substrate-specific adapters of a BCR (BTB-CUL3-RBX1) E3 ubiquitin-protein ligase complex involved in interferon response and anterograde Golgi to endosome transport. The BCR(KLHL20) E3 ubiquitin ligase complex mediates the ubiquitination of DAPK1, leading to its degradation by the proteasome, thereby acting as a negative regulator of apoptosis (Lee et al., 2010). Similarly to ankyrin-repeat, kelch-domain containing genes are located in variable genomic regions and show species-specific gene number and structure (Kochneva et al., 2005). Cowpox and sheeppox viruses lacking kelch proteins present virulence reduction (Lant & Maluquer de Motes, 2021). Kelch-protein A55, specifically, inhibits the NF-κB activity in response to IL-1β and TNF-α, as well as the transcription of NF-κB-responsive genes in response to TNF-α stimulation during Vaccinia virus infection (Pallett et al., 2019).Besides ankyrin and kelch-like proteins, genes related to DNA / RNA polymerase and helicase activity, early and late transcription factors, myristyl and core proteins, among others, were found to have sites under diversifying selective pressure. Interestingly, purifying selection pressure evidence was also observed, suggesting potentially important sites to molecular stability and functionality.

## 5 CONCLUSION

Analyzing the molecular evolution patterns of Monkeypox virus genomes, it is possible to observe some evidence of diversifying selection pressure on specific sites from protein coding sequences acting on immunomodulatory processes. The existence of different sites under diversifying - and purifying - selection in paralogs denotes adaptation mechanisms underlying the host-pathogen interaction of Monkeypox virus human species.

The presence of multiple nucleotide indels in MpxV genomes claims for posterior studies analyzing molecular evolution traits in these other regions, as well the detection of protein structural changes caused by the potential frame shifts and coding sequence truncations occurring in those genes. As a screening analysis, this study aims to identify potential targets involved in viral responses to the host immune system and their genetic alterations in front of new interaction pressures during pathogenesis and infection.

## DECLARATIONS

### Availability of data and materials

Full tables acknowledging the authors and corresponding labs submitting sequencing data used in this study can be found in Supplementary File 6. Additional information used and/or analyzed during the current study are available from the corresponding author on reasonable request.

### Competing interests

The authors declare no competing interests.

### Funding

This research did not receive any specific grant from funding agencies in the public, commercial, or not-for-profit sectors.

### CRediT statements

**Conceptualization:** Patrícia Aline Gröhs Ferrareze, Claudia Elizabeth Thompson; **Methodology:** Patrícia Aline Gröhs Ferrareze, Claudia Elizabeth Thompson**; Formal analysis and investigation:** Patrícia Aline Gröhs Ferrareze, Claudia Elizabeth Thompson**; Writing - original draft preparation:** Patrícia Aline Gröhs Ferrareze, Rute Alves Pereira e Costa, Claudia Elizabeth Thompson**; Writing - review and editing:** Patrícia Aline Gröhs Ferrareze, Rute Alves Pereira e Costa, Claudia Elizabeth Thompson**; Resources:** Claudia Elizabeth Thompson**; Supervision:** Claudia Elizabeth Thompson. All authors have read and approved the manuscript.

## Supplementary material

**Supplementary File 1**. Genomic alignment between Central African clade sequence DQ011155.1 (human Monkeypox virus strain Zaire_1979-005) and West African clade sequence NC_063383.1 (human Monkeypox virus strain M5312_HM12_Rivers). Identity shows synteny conservation between genomic blocks, which are colored in gray (syntenic) or black (non-syntenic). Annotated genes are colored in red and labeled. Predicted ORFs are colored in yellow. Genes with sites under positive selective pressure are highlighted in light red boxes.

**Supplementary File 2**. Gene identification correspondence from sequences with positively selected sites in orthogroups shared between Monkeypox, Vaccinia and Variola virus genomes.

**Supplementary File 3**. Synteny conservation between Central African clade sequence DQ011155.1 (human Monkeypox virus strain Zaire_1979-005) and West African clade sequence NC_063383.1 (human Monkeypox virus strain M5312_HM12_Rivers) for AAY97205 (A) and AAY97367 (B) sequence regions. Block colors indicate synteny conservation (pink), partial conservation (red) and synteny break (white). White horizontal bars indicate annotated genes. Blue arrows indicate predicted ortholog ORFs.

**Supplementary File 4**. Sites identified under positive and negative selective pressure by FUBAR and FEL methods, respectively, for Monkeypox virus genes and their associated biological processes.

**Supplementary File 5**. Maximum Likelihood phylogenetic tree from 155 concatenated Monkeypox virus gene alignments. (A) Partitioned tree. (B) Non-partitioned best-fit tree generated with TempEst software by evaluation of the sequence’s temporal signal.

**Supplementary File 6**. GISAID acknowledgment list for Monkeypox virus genomes selected for this study.

## Notes

### Competing Interest Statement

The authors have declared no competing interest.

