## Supplementary figures and images for "Genomic characterization and molecular evolution of human Monkeypox viruses"

### Supplementary File 1

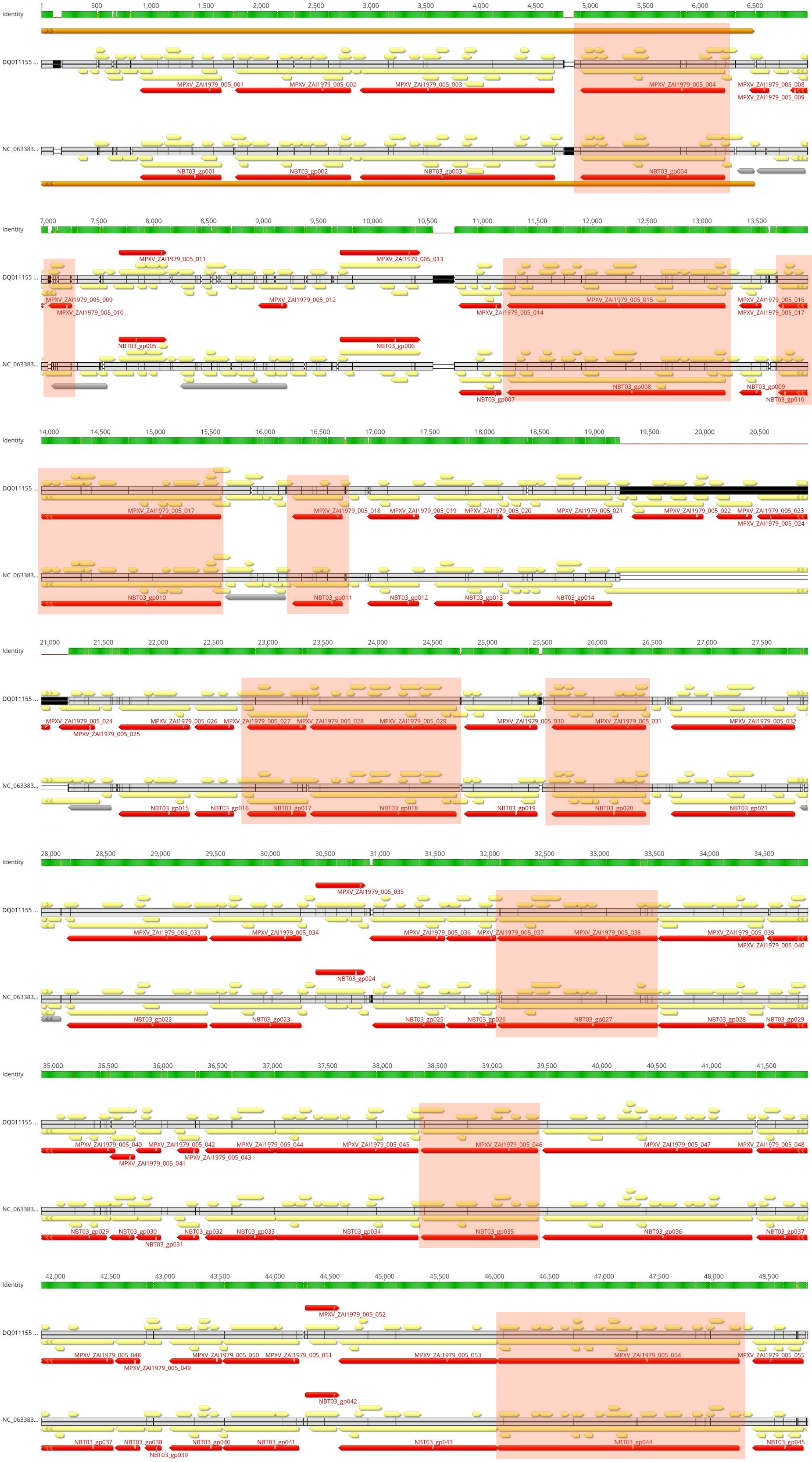

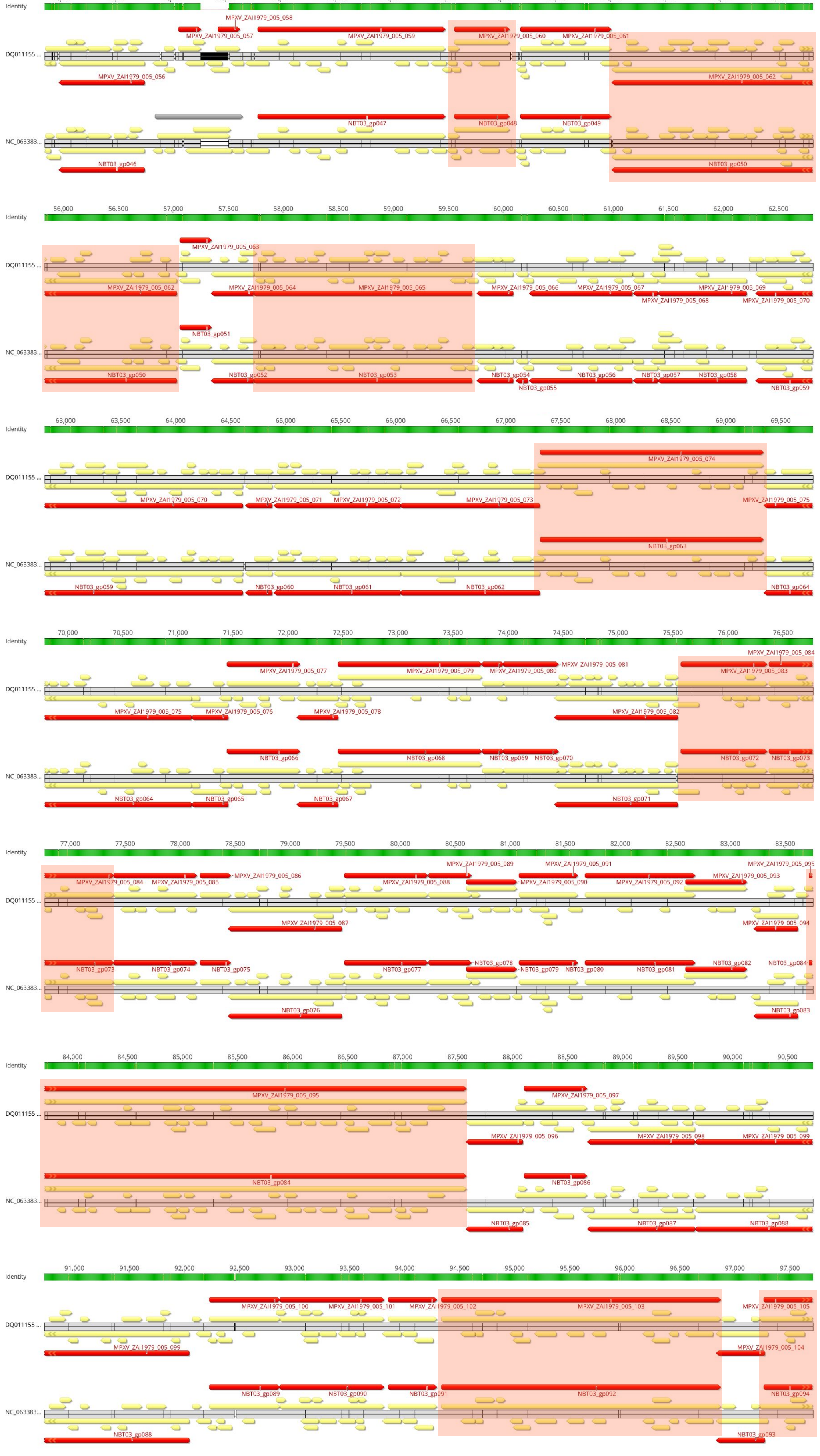

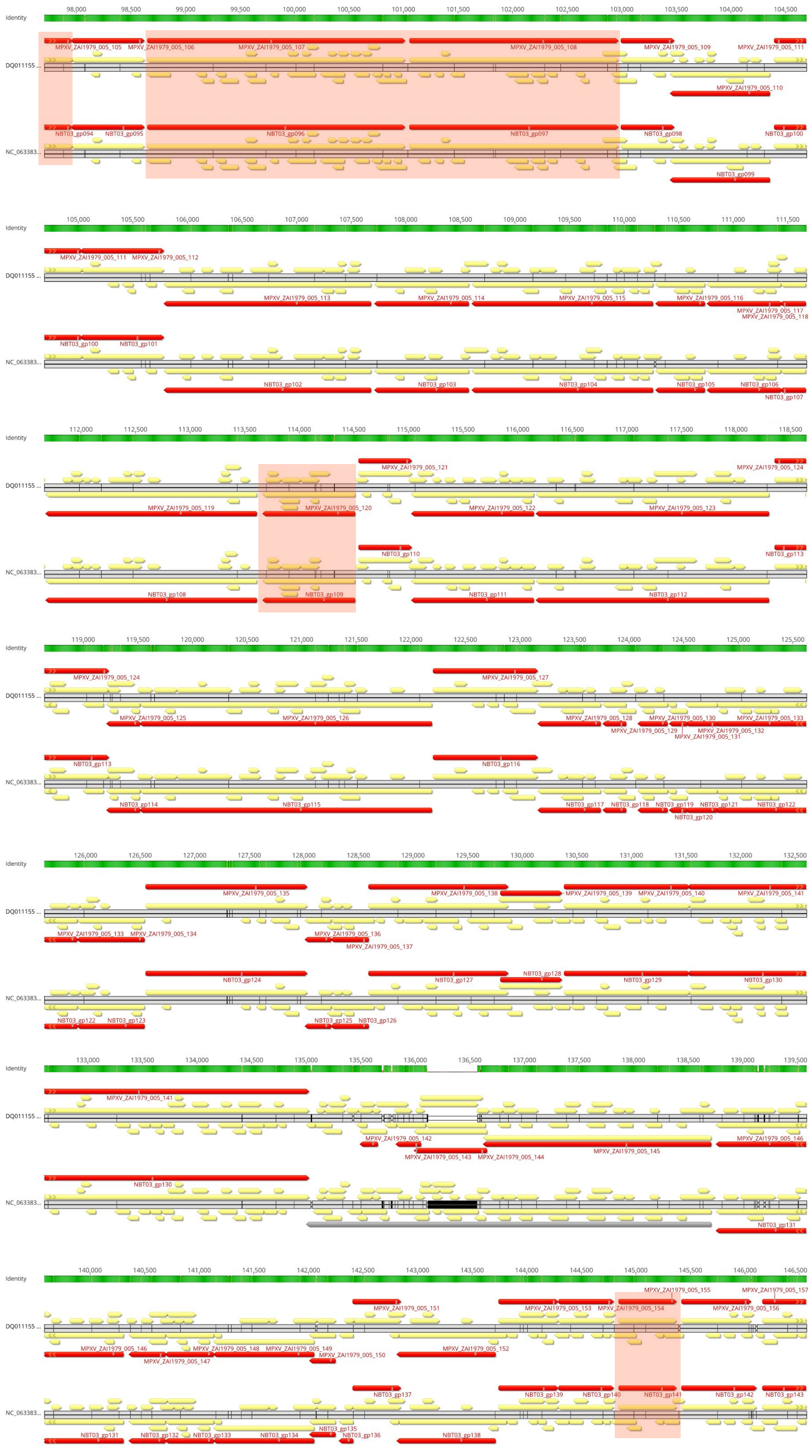

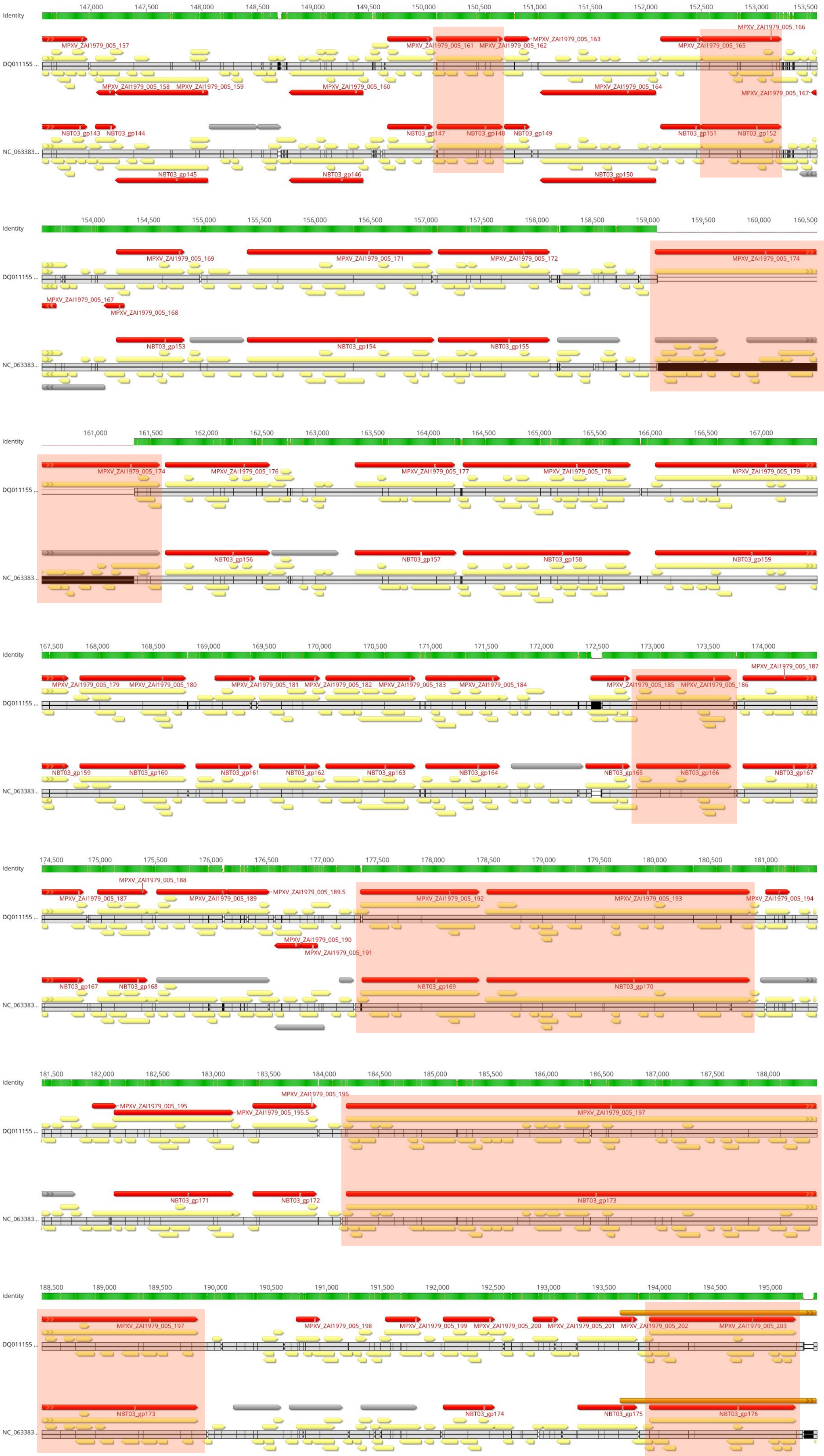

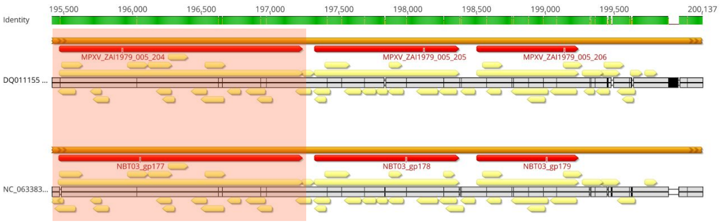

### Supplementary File 3

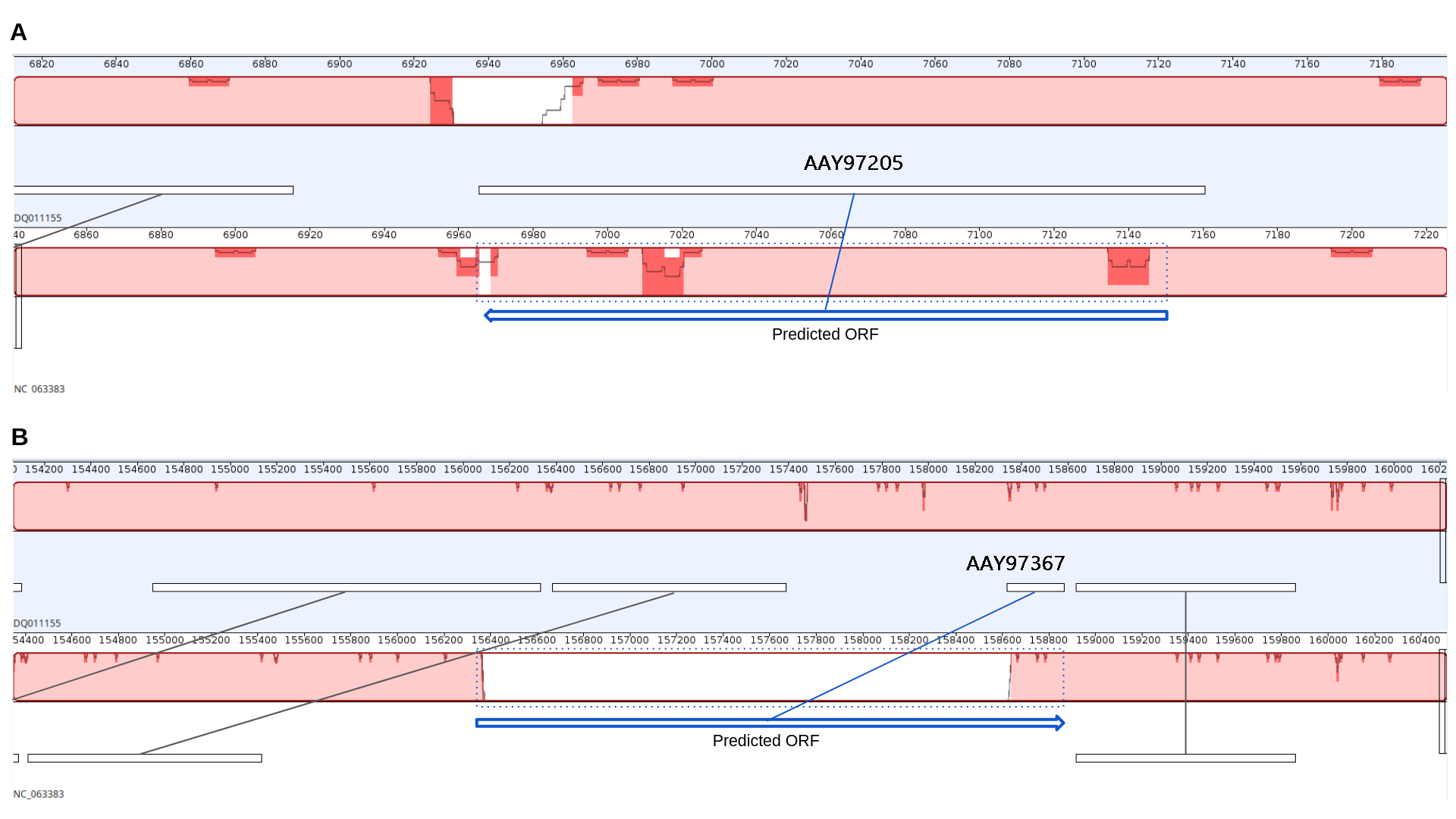

### Supplementary File 5

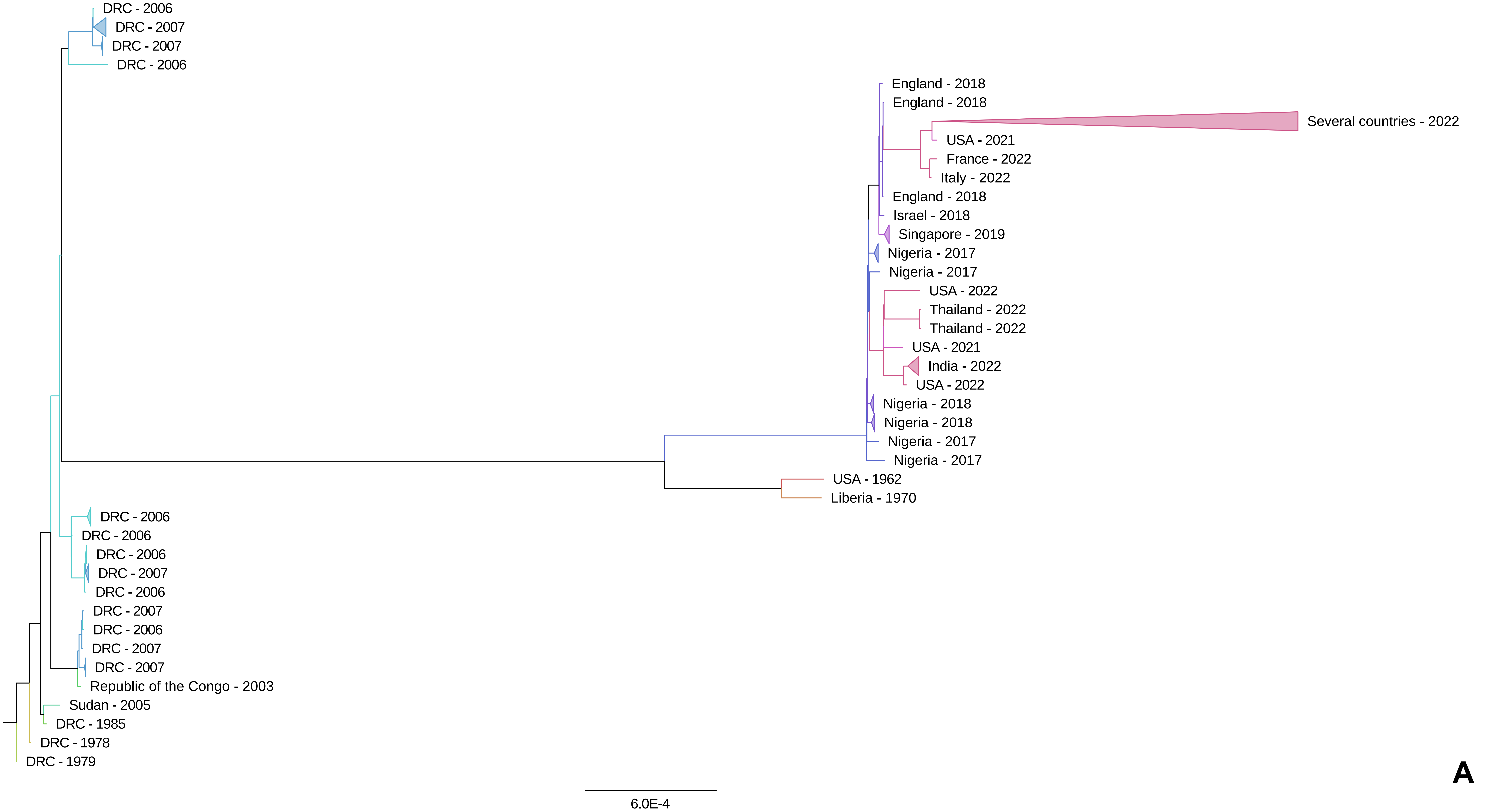

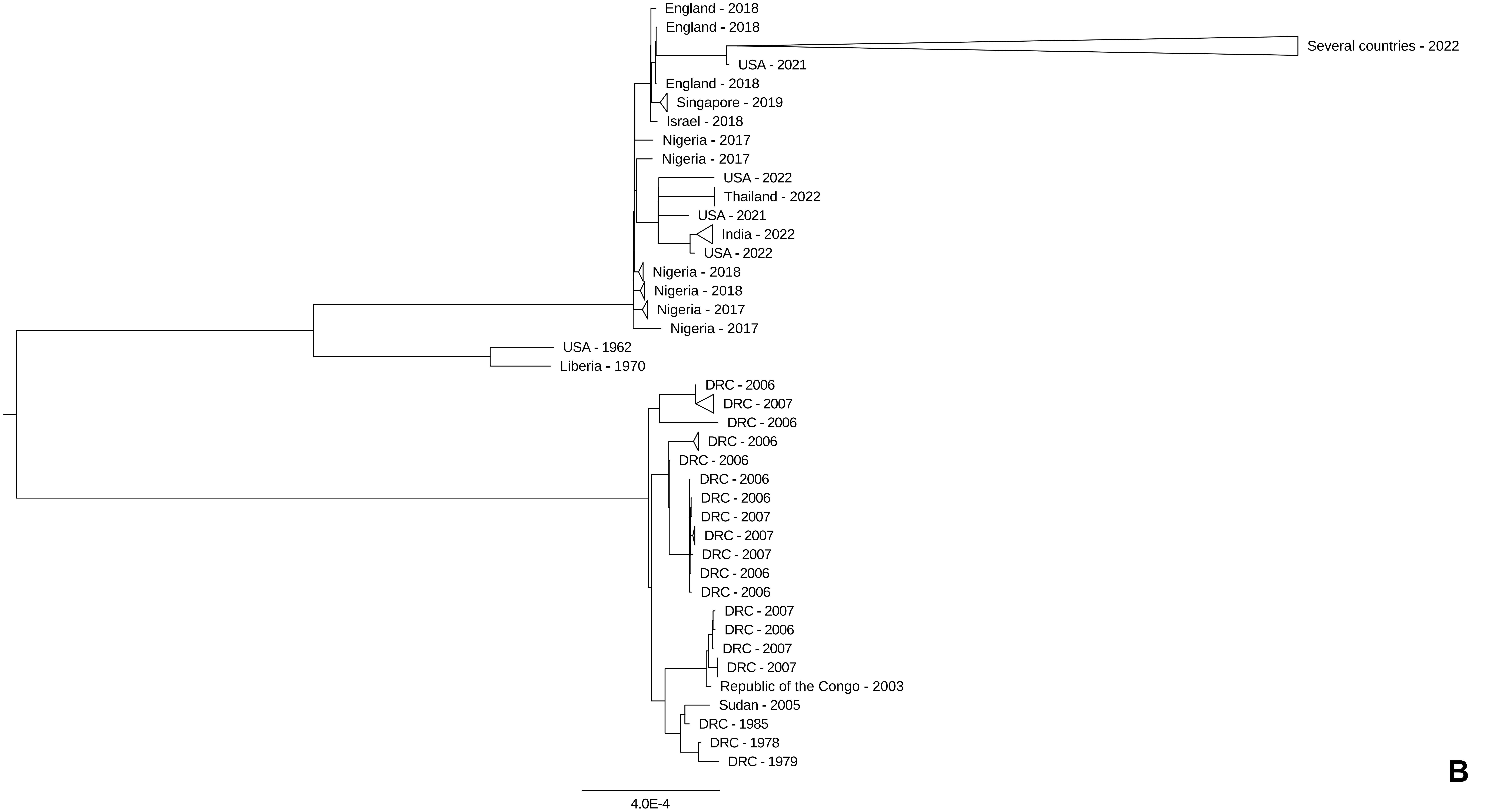
