## Supplementary File 2 for "Genomic characterization and molecular evolution of human Monkeypox viruses"

**Supplementary Table 2.** Orthogroups shared between Monkeypox, Vaccinia and Variola virus genomes.

| Monkeypox protein id | Protein name | Variola protein id | Vaccinia protein id |
|---|---|---|---|
| AAY97202 / AAY97397 | ankyrin-like protein | -- | -- |
| AAY97205 | unknown | -- | -- |
| AAY97210 | ankyrin-like protein | NP_042050 | QTC35413 / QTC35472 |
| AAY97212 | ankyrin-like protein | NP_042051 | QTC35512 / QTC35530 |
| AAY97213 | unknown | NP_042052 | QTC35398 |
| AAY97223 | alpha-amanitin target protein | NP_042061 | QTC35477 |
| AAY97224 | ankyrin-like protein | NP_042062 | -- |
| AAY97226 | ankyrin-like protein | NP_042064 / NP_042065 | QTC35503 |
| AAY97233 | kelch-like protein | NP_042071 | QTC35533 |
| AAY97241 | unknown | NP_042079 | QTC35442 |
| AAY97249 | unknown | NP_042087 | QTC35499 |
| AAY97255 | myristylprotein | NP_042092 | QTC35520 |
| AAY97257 | DNA polymerase | NP_042094 | QTC35382 |
| AAY97260 | unknown | NP_042097 | QTC35433 |
| AAY97269 | bifunctional DNA/RNA-helicase/DExH-NPH-II | NP_042106 | QTC35395 |
| AAY97278 | late gene transcription VLTF-1 | NP_042115 | QTC35496 |
| AAY97279 | myristylprotein | NP_042116 | QTC35521 |
| AAY97290 | DNA-dependent RNA polymerase subunit rpo147 | NP_042127 | QTC35488 |
| AAY97298 / AAY97299 | bifunctional large subunit of mRNA capping enzyme protein/transcription termination factor VTF | NP_042135 / NP_042136 | QTC35424 / QTC35445 |
| AAY97300 | virion core protein | NP_042137 | QTC35464 |
| AAY97302 | NTPase | NP_042139 | QTC35481 |
| AAY97303 | 70kDa small subunit of early gene transcription factor VETF | NP_042140 | QTC35370 |
| AAY97315 | 39kDa core protein | NP_042152 | QTC35375 |

| | | | |
|---|---|---|---|
| AAY97350 | unknown | NP_042186 | QTC35529 |
| AAY97357 | putative type-I membrane glycoprotein | NP_042198 | QTC35540 |
| AAY97361 | Toll/IL1-receptor [TIR]-like protein | NP_042202 | QTC35524 |
| AAY97367 | kelch-like protein | NP_042210 | -- |
| AAY97378 | ser/thr protein kinase-like protein | -- | QTC35421 |
| AAY97385 | IFN-alpha/beta-receptor-like secreted glycoprotein | NP_042232 | QTC35390 |
| AAY97386 | ankyrin-like protein | NP_042233 | -- |
| AAY97391 | putative membrane-associated glycoprotein | NP_042238 | |
| AAY97398 / AAY97201 | ankyrin-like protein | NP_042239 | -- |

Sets of two sequences separated by a "/" indicate paralog groups where the true orthologs were not defined by this analysis.
