## Supplementary File 6 for "Genomic characterization and molecular evolution of human Monkeypox viruses"

We gratefully acknowledge the following Authors from the Originating laboratories responsible for obtaining the specimens, as well as the Submitting laboratories where the genome data were generated and shared via GISAID, on which this research is based.

All Submitters of data may be contacted directly via www.gisaid.org

Authors are sorted alphabetically.

| Accession ID | Originating Laboratory | Submitting Laboratory | Authors |
|---|---|---|---|
| EPI_ISL_13052263 | Microbiol Genomics and Bioinformatics, Bundeswehr Institute of Microbiology | Microbiol Genomics and Bioinformatics, Bundeswehr Institute of Microbiology | Antwerpen,M.H., Lang,D., Zange,S., Walter,M.C. and Woelfel,R. |
| EPI_ISL_13052264, EPI_ISL_13052265, EPI_ISL_13052266, EPI_ISL_13052267, EPI_ISL_13052268, EPI_ISL_13052269, EPI_ISL_13052270, EPI_ISL_13052271, EPI_ISL_13052272, EPI_ISL_13052273 | Instituto Nacional de Saude Doutor Ricardo Jorge (INSA) | Instituto Nacional de Saude Doutor Ricardo Jorge (INSA) | Joana Isidro, Vítor Borges, Miguel Pinto, Daniel Sobral, João Dourado Santos, Alexandra Nunes, Verónica Mixão, Rita Ferreira, Daniela Santos, Silvia Duarte, Luís Vieira, Maria José Borrego, Sofia Núncio, Isabel Lopes de Carvalho, Ana Pelerito, Rita Cordeiro, João Paulo Gomes |
| EPI_ISL_13052274 | Laboratory of Virology, University Hospitals of Geneva | Laboratory of Virology, University Hospitals of Geneva | Laubscher,F., Chudzinski,V., Schibler,M., Kaiser,L. and Renzoni,A. |
| EPI_ISL_13052275 | IHAP, VIRAL, Universite de Toulouse, INRAE, ENVT | IHAP, VIRAL, Universite de Toulouse, INRAE, ENVT | Croville,G., Walch,M., Guerin,J.-L., Mansuy,J.-M., Pasquier,C. and Izopet,J. |
| EPI_ISL_13052277 | Public Health Virology, Erasmus Medical Centre | Public Health Virology, Erasmus Medical Centre | Oude Munnink,B.B., Boter,M., Wellers,B., Molenkamp,R., Sikkema,R.S. and Koopmans,M. |
| EPI_ISL_13052278 | Research and Evaluation, UKHSA | Research and Evaluation, UKHSA | Osman,K.L., Lewandowski,K.S., Pullan,S.T., Carter,D.P., Crook,J.M., Vipond,R. and Chand,M. |
| EPI_ISL_13052279, EPI_ISL_13052280, EPI_ISL_13052281 | Research and Evaluation, UKHSA | Research and Evaluation, UKHSA | Osman,K.L., Lewandowski,K.S., Carter,D.P., Crook,J.M., Pullan,S.T., Vipond,R. and Chand,M. |
| EPI_ISL_13052282 | Microbiology, Immunology and Transplantation, KU Leuven, Rega Institute | Microbiology, Immunology and Transplantation, KU Leuven, Rega Institute | Vanmechelen,B., Wawina-Bokalanga,T., Logist,A.-S., Sinnesael,R., Ysebaert,L., Verlinden,J., Bloemen,M. and Maes,P. |
| EPI_ISL_13052283 | Microbiology, Immunology and Transplantation, KU Leuven, Rega Institute | Microbiology, Immunology and Transplantation, KU Leuven, Rega Institute | Wawina-Bokalanga,T., Vanmechelen,B., Logist,A.-S., Sinnesael,R., Ysebaert,L., Verlinden,J., Bloemen,M. and Maes,P. |
| EPI_ISL_13052284 | Microbiology, Hospital Universitari Germans Trias i Pujol | Microbiology, Hospital Universitari Germans Trias i Pujol | Martinez-Puchol,S., Coello,A., Bordoy,A.E., Soler,L., Panisello,D., Gonzalez-Gomez,S., Clara,G., Paris de Leon,A., Not,A., Hernandez,A., Bofill-Mas,S., Saludes,V., Blanco,I., Martro,E. and Cardona,P.-J. |
| EPI_ISL_13052285 | Laboratory of Virology, University Hospitals of Geneva | Laboratory of Virology, University Hospitals of Geneva | Laubscher,F., Schibler,M., Kaiser,L. and Renzoni,A. |
| EPI_ISL_13052286 | Department of Biomedical and Clinical Sciences, University of Milan | Department of Biomedical and Clinical Sciences, University of Milan | Lai,A., Bergna,A., Della Ventura,C., Tarkowski,M., Riva,A., Moschese,D., Rizzardini,G., Antinori,S. and Zehender,G. |
| EPI_ISL_13052287 | Virology, GENomique EPIdemiologique des maladies Infectieuses | Virology, GENomique EPIdemiologique des maladies Infectieuses | unknown |
| EPI_ISL_13052288 | Department of Health, Utah Public Health Laboratory | Department of Health, Utah Public Health Laboratory | Young,E.L., Hergert,J. and Oakeson,K.F. |
| EPI_ISL_13052289 | Centers for Disease Control & Prevention (CDC), Division of High Consequence Pathogens and Pathology (DHCPP-PRB) | Centers for Disease Control & Prevention (CDC), Division of High Consequence Pathogens and Pathology (DHCPP-PRB) | Gigante,C.M., Smole,S., Seabolt,M.H., Wilkins,K., McCollum,A., Hutson,C., Davidson,W., Rao,A., Brown,C. and Li,Y. |
| EPI_ISL_13052290 | Laboratory for Diagnostics of Zoonoses and WHO Centre, Institute of Microbiology and Immunology, Faculty of Medicine, University of Ljubljana | Laboratory for Diagnostics of Zoonoses and WHO Centre, Institute of Microbiology and Immunology, Faculty of Medicine, University of Ljubljana | Zakotnik,S., Vlaj,D., Suljic,A., Zorec,T.M., Korva,M., Poljak,M. and Avsic Zupanc,T. |
| EPI_ISL_13052291 | Laboratory for Diagnostics of Zoonoses and WHO Centre, Institute of Microbiology and Immunology, Faculty of Medicine, University of Ljubljana | Laboratory for Diagnostics of Zoonoses and WHO Centre, Institute of Microbiology and Immunology, Faculty of Medicine, University of Ljubljana | Zakotnik,S., Vlaj,D., Suljic,A., Zorec,T.M., Skubic,C., Rozman,D., Korva,M., Poljak,M. and Avsic Zupanc,T. |
| EPI_ISL_13052292 | Victorian Infectious Diseases Reference Laboratory, Doherty Institute | Victorian Infectious Diseases Reference Laboratory, Doherty Institute | Hammerschlag,Y., MacLeod,G., Papadakis,G., Adan-Sanchez,A., Druce,J.D., Williamson,D.A., Cheng,A.C. and McMahon,J.H. |
| EPI_ISL_13052293, EPI_ISL_13052294 | Centre for Biological Threats, Highly Pathogenic Viruses, Robert Koch Institute | Centre for Biological Threats, Highly Pathogenic Viruses, Robert Koch Institute | Brinkmann,A., Kohl,C., Uddin,S., Pape,K., Schrick,L., Michel,J., Schaade,L. and Nitsche,A. |
| EPI_ISL_13052295 | SC (UCO) Igiene e Sanità Pubblica, ASUGI, Trieste | Genomics and Epigenomics, AREA Science Park | Licastro,D., DeGasperi,M., Negri,C., Piscianz,E., Koncan,R., Dal Monego,S., Segat,L. and D'Agaro,P. |
| EPI_ISL_13053218 | National Center for Infectious Diseases, Centers for Disease Control and Prevention | National Center for Infectious Diseases, Centers for Disease Control and Prevention | Likos,A.M., Sammons,S.A., Olson,V.A., Frace,A.M., Li,Y., Olsen-Rasmussen,M., Davidson,W., Galloway,R., Khristova,M.L., Reynolds,M.G., Zhao,H., Carroll,D.S., Curns,A., Formenty,P., Esposito,J.J., Regnery,R.L. and Damon,I.K. |
| EPI_ISL_13056233, EPI_ISL_13056234, EPI_ISL_13056235, EPI_ISL_13056236, EPI_ISL_13056237, EPI_ISL_13056238, EPI_ISL_13056239, EPI_ISL_13056240, EPI_ISL_13056241, EPI_ISL_13056242, EPI_ISL_13056243, EPI_ISL_13056244, EPI_ISL_13056245, EPI_ISL_13056246, EPI_ISL_13056247, EPI_ISL_13056248, EPI_ISL_13056249, EPI_ISL_13056250, EPI_ISL_13056251, EPI_ISL_13056252, EPI_ISL_13056253, EPI_ISL_13056254, EPI_ISL_13056255 | | | |
| see above | USAMRIID, Center for Genome Sciences, United States Army Medical Research Institute of Infectious Diseases | USAMRIID, Center for Genome Sciences, United States Army Medical Research Institute of Infectious Diseases | Kugelman,J.R., Johnston,S.C., Mulembakani,P.M., Kisalu,N., Lee,M.S., Koroleva,G., McCarthy,S.E., Gestole,M.C., Wolfe,N.D., Fair,J.N., Schneider,B.S., Wright,L.L., Huggins,J., Whitehouse,C.A., Wemakoy,E.O., Muyembe-Tamfum,J.J., Hensley,L.E., Palacios,G.F. and Rimoin,A.W. |
| EPI_ISL_13056271, EPI_ISL_13056272, EPI_ISL_13056273, EPI_ISL_13056274 | Centers for Disease Control and Prevention | Centers for Disease Control and Prevention | Mauldin,M.R., McCollum,A.M., Nakazawa,Y.J., Mandra,A., Whitehouse,E.R., Davidson,W., Zhao,H., Gao,J., Li,Y., Doty,J., Yinka-Ogunleye,A., Akinpelu,A., Aruna,O., Naidoo,D., Lewandowski,K., Afrough,B., Graham,V., Aarons,E., Hewson,R., Vipond,R., Dunning,J., Chand,M., Brown,C., Cohen-Gihon,I., Erez,N., Shifman,O., Israeli,O., Sharon,M., Schwartz,E., Beth-Din,A., Zvi,A., Mak,T.M., Ng,Y.K., Cui,L., Lin,R.T.P., Olson,V.A., Brooks,T., Paran,N., Ihekweazu,C. and Reynolds,M.G. |
| EPI_ISL_13056275, EPI_ISL_13056276, EPI_ISL_13056277, EPI_ISL_13056278, EPI_ISL_13056279, EPI_ISL_13056280, EPI_ISL_13056281 | Centers for Disease Control and Prevention | Centers for Disease Control and Prevention | Yinka-Ogunleye,A., Aruna,O., Dalhat,M., Ogoina,D., McCollum,A., Disu,Y., Mamadu,I., Akinpelu,A., Ahmad,A., Burga,J., Ndoreraho,A., Nkunzimana,E., Manneh,L., Mohammed,A., Adeoye,O., Tom-Aba,D., Silenou,B., Ipadeola,O., Saleh,M., Adeyemo,A., Nwadiutor,I., Aworabhi,N., Uke,P., John,D., Wakama,P., Reynolds,M., Mauldin,M., Doty,J., Wilkins,K., Musa,J., Khalakdina,A., Adedeji,A., Mba,N., Ojo,O., Krause,G. and Ihekweazu,C. |
| EPI_ISL_13056282, EPI_ISL_13056283, EPI_ISL_13056284, EPI_ISL_13056285 | Centers for Disease Control and Prevention | Centers for Disease Control and Prevention | Mauldin,M.R., McCollum,A.M., Nakazawa,Y.J., Mandra,A., Whitehouse,E.R., Davidson,W., Zhao,H., Gao,J., Li,Y., Doty,J., Yinka-Ogunleye,A., Akinpelu,A., Aruna,O., Naidoo,D., Lewandowski,K., Afrough,B., Graham,V., Aarons,E., Hewson,R., Vipond,R., Dunning,J., Chand,M., Brown,C., Cohen-Gihon,I., Erez,N., Shifman,O., Israeli,O., Sharon,M., Schwartz,E., Beth-Din,A., Zvi,A., Mak,T.M., Ng,Y.K., Cui,L., Lin,R.T.P., Olson,V.A., Brooks,T., Paran,N., Ihekweazu,C. and Reynolds,M.G. |
| EPI_ISL_13056289 | Biochemistry and Molecular Biology, Israel Institute for Biological Research | Biochemistry and Molecular Biology, Israel Institute for Biological Research | Cohen Gihon,I., Israeli,O., Shifman,O., Erez,N., Melamed,S., Paran,N., Beth-Din,A. and Zvi,A. |
| EPI_ISL_13056556 | Walter Reed Army Institute of Research | Biochemistry and Microbiology, University of Victoria | Chen,N., Li,G., Liszewski,M.K., Atkinson,J.P., Jahrling,P.B., Feng,Z., Schriewer,J., Buck,C., Wang,C., Lefkowitz,E.J., Esposito,J.J., Harms,T., Damon,I.K., Roper,R.L., Upton,C. and Buller,R.M. |
| EPI_ISL_13056892, EPI_ISL_13056893, EPI_ISL_13056894, EPI_ISL_13056895, EPI_ISL_13056896, EPI_ISL_13056897, EPI_ISL_13056898, EPI_ISL_13056899, EPI_ISL_13056900, EPI_ISL_13056901, EPI_ISL_13056902, EPI_ISL_13056903, EPI_ISL_13056904, EPI_ISL_13056905, EPI_ISL_13056906, EPI_ISL_13056907, | | | |

| | | | |
|---|---|---|---|
| EPI_ISL_13056908, EPI_ISL_13056909 | | | |
| see above | Instituto Nacional de Saude Doutor Ricardo Jorge (INSA) | Instituto Nacional de Saude Doutor Ricardo Jorge (INSA) | Joana Isidro, Vítor Borges, Miguel Pinto, Daniel Sobral, João Dourado Santos, Alexandra Nunes, Verónica Mixão, Rita Ferreira, Daniela Santos, Silvia Duarte, Luís Vieira, Maria José Borrego, Sofia Núncio, Isabel Lopes de Carvalho, Ana Pelerito, Rita Cordeiro, João Paulo Gomes |
| EPI_ISL_13056910 | Biochemistry and Molecular Genetics, Israel Institute for Biological Research | Biochemistry and Molecular Genetics, Israel Institute for Biological Research | Israeli,O., Guedj-Dana,Y., Lazar,S., Shifman,O., Erez,N., Weiss,S., Paran,N., Israely,T., Schuster,O., Zvi,A., Beth-Din,A. and Cohen Gihon,I. |
| EPI_ISL_13058404, EPI_ISL_13058405 | National Center for Infectious Diseases, Centers for Disease Control and Prevention | National Center for Infectious Diseases, Centers for Disease Control and Prevention | Likos,A.M., Sammons,S.A., Olson,V.A., Frace,A.M., Li,Y., Olsen-Rasmussen,M., Davidson,W., Galloway,R., Khristova,M.L., Reynolds,M.G., Zhao,H., Carroll,D.S., Curns,A., Formenty,P., Esposito,J.J., Regnery,R.L. and Damon,I.K. |
| EPI_ISL_13058456 | Vaccine and Gene Therapy Institute, Oregon Health and Science University | Vaccine and Gene Therapy Institute, Oregon Health and Science University | Estep,R.D., Messaoudi,I., O'Connor,M.A., Li,H., Sprague,J., Barron,A., Engelmann,F., Yen,B., Powers,M.F., Jones,J.M., Robinson,B.A., Orzechowska,B.U., Manoharan,M., Legasse,A., Planer,S., Wilk,J., Axthelm,M.K. and Wong,S.W. |
| EPI_ISL_13058459, EPI_ISL_13058460 | Biochemistry and Microbiology, University of Victoria | Biochemistry and Microbiology, University of Victoria | Nakazawa,Y., Emerson,G.L., Carroll,D.S., Zhao,H., Li,Y., Reynolds,M.G., Karem,K.L., Olson,V.A., Lash,R.R., Davidson,W.B., Smith,S.K., Levine,R.S., Regnery,R.L., Sammons,S.A., Frace,M.A., Mutasim,E.M., Karsani,M.E., Muntasir,M.O., Babiker,A.A., Opoka,L., Chowdhary,V. and Damon,I.K. |
| EPI_ISL_13058475 | National Public Health Laboratory, National Centre for Infectious Diseases | National Public Health Laboratory, National Centre for Infectious Diseases | Yong,S.E.F., Ng,O.T., Ho,Z.J.M., Mak,T.M., Marimuthu,K., Vasoo,S., Yeo,T.W., Ng,Y.K., Cui,L., Ferdous,Z., Chia,P.Y., Aw,B.J.W., Manauis,C.M., Low,C.K.K., Chan,G., Peh,X., Lim,P.L., Chow,L.P.A., Chan,M., Lee,V.J.M., Lin,R.T.P., Heng,M.K.D. and Leo,Y.S. |
| EPI_ISL_13089461 | Hospital General Universitario Gregorio Marañón | Hospital General Universitario Gregorio Marañón | Sergio Buenestado Serrano, Rosalía Palomino Cabrera, Daniel Peñas Utrilla, Jorge Rodríguez-Grande, Laura Pérez-Lago, Cristina Rodriguez-Grande, Marta Herranz Martin, Julia Suárez, Pilar Catalán, Patricia Muñoz, Darío García de Viedma |
| EPI_ISL_13094227 | Centers for Disease Control & Prevention (CDC), Division of High Consequence Pathogens and Pathology (DHCPP-PRB) | Centers for Disease Control & Prevention (CDC), Division of High Consequence Pathogens and Pathology (DHCPP-PRB) | Gigante,C.M., Lee,P., Seabolt,M.H., Wilkins,K., McCollum,A., Hutson,C., Davidson,W., Rao,A., Mendoza,R. and Li,Y. |
| EPI_ISL_13096615 | Centers for Disease Control & Prevention (CDC), Division of High Consequence Pathogens and Pathology (DHCPP-PRB) | Centers for Disease Control & Prevention (CDC), Division of High Consequence Pathogens and Pathology (DHCPP-PRB) | Gigante,C.M., Griffin-Thomas,L.A., Seabolt,M.H., Wilkins,K., McCollum,A., Hutson,C., Davidson,W., Rao,A., Crain,J. and Li,Y. |
| EPI_ISL_13100618 | Centers for Disease Control & Prevention (CDC), Division of High Consequence Pathogens and Pathology (DHCPP-PRB) | Centers for Disease Control & Prevention (CDC), Division of High Consequence Pathogens and Pathology (DHCPP-PRB) | Gigante,C.M., Ventura,J., Seabolt,M.H., Wilkins,K., McCollum,A., Hutson,C., Davidson,W., Rao,A., Nash,J. and Li,Y. |
| EPI_ISL_13100619 | Centers for Disease Control & Prevention (CDC), Division of High Consequence Pathogens and Pathology (DHCPP-PRB) | Centers for Disease Control & Prevention (CDC), Division of High Consequence Pathogens and Pathology (DHCPP-PRB) | Gigante,C.M., Lee,P., Seabolt,M.H., Wilkins,K., McCollum,A., Hutson,C., Davidson,W., Rao,A., Mendoza,R. and Li,Y. |
| EPI_ISL_13100620 | Centers for Disease Control & Prevention (CDC), Division of High Consequence Pathogens and Pathology (DHCPP-PRB) | Centers for Disease Control & Prevention (CDC), Division of High Consequence Pathogens and Pathology (DHCPP-PRB) | Gigante,C.M., Atkinson,A., Seabolt,M.H., Wilkins,K., McCollum,A., Hutson,C., Davidson,W., Rao,A., Murray,J. and Li,Y. |
| EPI_ISL_13100621 | Centers for Disease Control & Prevention (CDC), Division of High Consequence Pathogens and Pathology (DHCPP-PRB) | Centers for Disease Control & Prevention (CDC), Division of High Consequence Pathogens and Pathology (DHCPP-PRB) | Gigante,C.M., Stringer,J., Seabolt,M.H., Wilkins,K., McCollum,A., Hutson,C., Davidson,W., Rao,A., Schulte,J. and Li,Y. |
| EPI_ISL_13100622 | Centers for Disease Control & Prevention (CDC), Division of High Consequence Pathogens and Pathology (DHCPP-PRB) | Centers for Disease Control & Prevention (CDC), Division of High Consequence Pathogens and Pathology (DHCPP-PRB) | Gigante,C.M., Myers,R., Seabolt,M.H., Wilkins,K., McCollum,A., Hutson,C., Davidson,W., Rao,A., Blythe,D. and Li,Y. |
| EPI_ISL_13100719 | Centers for Disease Control & Prevention (CDC), Division of High Consequence Pathogens and Pathology (DHCPP-PRB) | Centers for Disease Control & Prevention (CDC), Division of High Consequence Pathogens and Pathology (DHCPP-PRB) | Gigante,C.M., Ventura,J., Seabolt,M.H., Wilkins,K., McCollum,A., Hutson,C., Davidson,W., Rao,A., Nash,J. and Li,Y. |
| EPI_ISL_13106454 | Hospital General Universitario Gregorio Marañón | Hospital General Universitario Gregorio Marañón | Sergio Buenestado Serrano, Rosalía Palomino Cabrera, Daniel Peñas Utrilla, Jorge Rodríguez-Grande, Pedro Sola Campoy, Laura Pérez-Lago, Cristina Rodriguez-Grande, Marta Herranz Martin, Julia Suárez, Pilar Catalán, Patricia Muñoz, Darío García de Viedma |
| EPI_ISL_13117291 | Centre for Biological Threats, Highly Pathogenic Viruses, Robert Koch Institute | Centre for Biological Threats, Highly Pathogenic Viruses, Robert Koch Institute | Brinkmann,A., Kohl,C., Uddin,S., Pape,K., Schrick,L., Michel,J., Jessen,H., Schaade,L. and Michel,A. |
| EPI_ISL_13117292, EPI_ISL_13117293 | Centre for Biological Threats, Highly Pathogenic Viruses, Robert Koch Institute | Centre for Biological Threats, Highly Pathogenic Viruses, Robert Koch Institute | Brinkmann,A., Kohl,C., Uddin,S., Pape,K., Schrick,L., Michel,J., Stocker,H., Schaade,L. and Nitsche,A. |
| EPI_ISL_13117294, EPI_ISL_13117295, EPI_ISL_13117296, EPI_ISL_13117297, EPI_ISL_13117298 | Centre for Biological Threats, Highly Pathogenic Viruses, Robert Koch Institute | Centre for Biological Threats, Highly Pathogenic Viruses, Robert Koch Institute | Brinkmann,A., Kohl,C., Uddin,S., Pape,K., Schrick,L., Michel,J., Schaade,L. and Nitsche,A. |
| EPI_ISL_13148263, EPI_ISL_13148264, EPI_ISL_13148265 | Centre for Biological Threats, Highly Pathogenic Viruses, Robert Koch Institute | Centre for Biological Threats, Highly Pathogenic Viruses, Robert Koch Institute | Brinkmann,A., Kohl,C., Uddin,S., Pape,K., Schrick,L., Michel,J., Jessen,H., Schaade,L. and Nitsche,A. |
| EPI_ISL_13148266, EPI_ISL_13148267, EPI_ISL_13148268, EPI_ISL_13148269 | Centre for Biological Threats, Highly Pathogenic Viruses, Robert Koch Institute | Centre for Biological Threats, Highly Pathogenic Viruses, Robert Koch Institute | Brinkmann,A., Kohl,C., Uddin,S., Pape,K., Schrick,L., Michel,J., Stocker,H., Schaade,L. and Nitsche,A. |
| EPI_ISL_13148270, EPI_ISL_13148271, EPI_ISL_13148272, EPI_ISL_13148273, EPI_ISL_13148274, EPI_ISL_13148275, EPI_ISL_13148276 | Centre for Biological Threats, Highly Pathogenic Viruses, Robert Koch Institute | Centre for Biological Threats, Highly Pathogenic Viruses, Robert Koch Institute | Brinkmann,A., Kohl,C., Uddin,S., Pape,K., Schrick,L., Michel,J., Schaade,L. and Nitsche,A. |
| EPI_ISL_13191438 | Instituto de Infectologia Emilio Ribas | Instituto Adolfo Lutz Strategic Laboratory | Claudio Tavares Sacchi, Karoline Rodrigues Campos, Marlon Benedito Nascimento Santos, Alex Domingos Reis, Ariadne Ferreira Amarante, Adriano Abbud, Adriana Bugno, Walkiria Delnoro Almeida Prado, Regiane Cardoso de Paula |
| EPI_ISL_13194516 | Alberta Precision Laboratories | Alberta Precision Laboratories | Matthew Croxen, Ashwin Deo, Paul Dieu, Xiaoli Dong, Kara Gill, David Granger, Christina Ferrato, Vanipriyadarsini Ikkurti, Jamil Kanji, Petya Koleva, Vincent Li, Colin Lloyd, Tarah Lynch, Raymond Ma, Kanti Pabbaraju, Silas Rotich, Hilary Sergeant, Steven Shideler, Todd Skitsko, Sandy Shokopoles, Graham Tipples, Johanna Thayer, Anita Wong |
| EPI_ISL_13234112 | Laboratório Central de Saúde Pública do Estado do Rio Grande do Sul | Instituto Adolfo Lutz Strategic Laboratory | Claudio Tavares Sacchi, Karoline Rodrigues Campos, Adriano Abbud, Adriana Bugno |
| EPI_ISL_13242738 | Hospital General Universitario Gregorio Marañón | Hospital General Universitario Gregorio Marañón | Sergio Buenestado Serrano, Rosalía Palomino Cabrera, Daniel Peñas Utrilla, Jorge Rodríguez-Grande, Pedro Sola Campoy, Laura Pérez-Lago, Cristina Rodriguez-Grande, Marta Herranz Martin, Julia Suárez, Pilar Catalán, Patricia Muñoz, Darío García de Viedma |
| EPI_ISL_13244349 | Erasmus Medical Center Department of Virology | Erasmus Medical Center Department of Virology | Bas Oude Munnink, Marjan Boter, Babette Weller, Richard Molenkamp, Janette Rahamat-Langendoen, Reina Sikkema, Marion Koopmans |
| EPI_ISL_13251120 | Laboratory of Virology, INMI Lazzaro Spallanzani IRCCS | Laboratory of Virology, INMI Lazzaro Spallanzani IRCCS | Giombini,E., Gruber,C.E.M., Rueca,M., Gramigna,G., VIta,S., Carletti,F., D'Abramo,A., Lapa,D., Puro,V., Fabeni,L., Butera,O., Colavita,F., Meschi,S., Matusali,G., Specchiarello,E., Vairo,F., Vaia,F., Garbuglia,A.R., Nicastri,E., Antinori,A., GIrardi,E. and Maggi,F. |
| EPI_ISL_13251157 | checkin Zollhaus | Institute of Medical Virology, University of Zurich | Verena Kufner, Gabriela Ziltener, Maryam Zaheri, Stefan Schmutz, Annette Audigé, Odette Bernasconi, Kevin Steiner, Jon Huder, Cyril Shah, Riccarda Capaul, Guido Bloemberg, Jürg Böni, Michael Huber, Alexandra Trkola |
| EPI_ISL_13251584 | Division of Infectious Diseases, University Hospital Zürich | Institute of Medical Virology, University of Zurich | Verena Kufner, Gabriela Ziltener, Maryam Zaheri, Stefan Schmutz, Annette Audigé, Odette Bernasconi, Kevin Steiner, Jon Huder, Cyril Shah, Riccarda Capaul, Guido Bloemberg, Jürg Böni, Michael Huber, Alexandra Trkola |
| EPI_ISL_13251723 | checkin Zollhaus | Institute of Medical Virology, University of Zurich | Verena Kufner, Gabriela Ziltener, Maryam Zaheri, Stefan Schmutz, Annette Audigé, Odette Bernasconi, Kevin Steiner, Jon Huder, Cyril Shah, Riccarda Capaul, Guido Bloemberg, Jürg Böni, Michael Huber, Alexandra Trkola |
| EPI_ISL_13269478 | Alberta Precision Laboratories | Alberta Precision Laboratories | Matthew Croxen, Ashwin Deo, Paul Dieu, Xiaoli Dong, Kara Gill, David Granger, Christina Ferrato, Vanipriyadarsini Ikkurti, Jamil Kanji, Petya Koleva, Vincent Li, Colin Lloyd, Tarah Lynch, Raymond Ma, Kanti Pabbaraju, Silas Rotich, Hilary Sergeant, Steven Shideler, Todd Skitsko, Sandy Shokopoles, Graham Tipples, Johanna Thayer, Anita Wong |
| EPI_ISL_13270980 | Instituto de Infectologia Emilio Ribas | Imperial College London, School of Public Health | Claro,I.M., de Lima,E.L., Romano,C.M., Candido,D.S., Lindoso,J.A.L., Barra,L.A.C., Borges,L.M.S., Medeiros,L.A., Tomishige,M.Y.S., Ramundo,M.S., Moutinho,T., da Silva,A.J.D., Rodrigues,C.C.M., de Azevedo,L.C.F., Villas-Boas,L.S., da Silva,C.A.M., Coletti,T.M., O'Toole,A., Quick,J., Loman,N., |

| | | | |
|---|---|---|---|
| | | | Rambaut,A., Faria,N.R., Figueiredo-Mello,C. and Sabino,E.C. |
| EPI_ISL_13302316 | Laboratory of Clinical Microbiology, Virology and Bioemergencies. ASST-Fatebenefratelli-Sacco, L.Sacco University Hospital | Army Medical and Veterinary Research Center | Silvia Fillo, Riccardo De Sanctis, Giovanni Faggioni, Andrea Ciammaruconi, Anna Anselmo, Vanessa Vera Fain, Simone Di Sabatino, Francesco Giordani, Antonella Fortunato, Rossella Brandi, Giulia Campoli, Marzia Cavalli, Anella Monte, Martina Lipari, Maria Di Spirito, Giorgia Grilli, Silvia Chimienti, Giandomenico Cerreto, Filippo Molinari, Giancarlo Petralito, Davide Mileto, Valeria Micheli, Maria Rita Gismondo, Florigio Lista |
| EPI_ISL_13304977 | National Public Health Center, National Biosafety Laboratory | National Public Health Center, National Biosafety Laboratory | Judit Henczkó, Dániel Déri, Lili Jármi, Bernadett Pályi, Zoltán Kis, |
| EPI_ISL_13308117, EPI_ISL_13308118, EPI_ISL_13308119, EPI_ISL_13308121, EPI_ISL_13308122, EPI_ISL_13308124, EPI_ISL_13308125, EPI_ISL_13308127, EPI_ISL_13308129, EPI_ISL_13308131, EPI_ISL_13308133, EPI_ISL_13308135, EPI_ISL_13308137, EPI_ISL_13308139, EPI_ISL_13308140, EPI_ISL_13308142, EPI_ISL_13308144, EPI_ISL_13308145, EPI_ISL_13308146, EPI_ISL_13308147, EPI_ISL_13308148, EPI_ISL_13308150, EPI_ISL_13308151, EPI_ISL_13308153, EPI_ISL_13308155, EPI_ISL_13308157 | | | |
| see above | Centre for Biological Threats, Highly Pathogenic Viruses, Robert Koch Institute | Centre for Biological Threats, Highly Pathogenic Viruses, Robert Koch Institute | Brinkmann,A., Kohl,C., Uddin,S., Pape,K., Schrick,L., Michel,J., Schaade,L. and Nitsche,A. |
| EPI_ISL_13308158, EPI_ISL_13308160 | IRBA Research Institute Biomédicale Des Armées | IRBA Research Institute Biomédicale Des Armées | Jarjaval,F., Nolent,F., Criqui,A., Chapus,C., Lamer,O., Ferraris,O. and Gorge,O. |
| EPI_ISL_13308162, EPI_ISL_13308163, EPI_ISL_13308165, EPI_ISL_13308167 | Laboratory for Diagnostics of Zoonoses and WHO Centre, Institute of Microbiology and Immunology, Faculty of Medicine, University of Ljubljana | Laboratory for Diagnostics of Zoonoses and WHO Centre, Institute of Microbiology and Immunology, Faculty of Medicine, University of Ljubljana | Zakotnik,S., Vlaj,D., Suljic,A., Zorec,T.M., Korva,M., Poljak,M. and Avsic Zupanc,T. |
| EPI_ISL_13314740 | Laboratorio de Vigilancia em Saude de Vinhedo | Instituto Adolfo Lutz Strategic Laboratory | Claudio Tavares Sacchi, Karoline Rodrigues Campos, Adriano Abbud, Adriana Bugno |
| EPI_ISL_13331598 | Department for Virology, Molecular Biology and Genome Research, R. G. Lugar Center for Public Health Research, National Center for Disease Control and Public Health (NCDC) of Georgia | Department for Virology, Molecular Biology and Genome Research, R. G. Lugar Center for Public Health Research, National Center for Disease Control and Public Health (NCDC) of Georgia | Giorgi Tomashvili, Salome Javashvili, Meri Pantsulaia, Gvantsa Brachveli, Ana Papkiauri, Gvantsa Chanturia, Adam Kotorashvili, Maia Alkhazashvili, Khatuna Zakhashvili, Paata Imnadze, Amiran Gamkrelidze. |
| EPI_ISL_13331712 | Laboratory of Virology, INMI Lazzaro Spallanzani IRCCS | Laboratory of Virology, INMI Lazzaro Spallanzani IRCCS | Rueca,M., Giombini,E., Gruber,C.E.M., Gramigna,G., Mazzotta,V., Carletti,F., Lapa,D., Pittalis,S., Puro,V., Fabeni,L., Butera,O., Colavita,F., Meschi,S., Matusali,G., Specchiarello,E., Vairo,F., Vaia,F., Nicastri,E., Antinori,A., Girardi,E. and Maggi,F. |
| EPI_ISL_13331713 | Laboratory of Virology, INMI Lazzaro Spallanzani IRCCS | Laboratory of Virology, INMI Lazzaro Spallanzani IRCCS | Gramigna,G., Giombini,E., Gruber,C.E.M., Rueca,M., Carletti,F., Cicalini,S., Lapa,D., Puro,V., Marani,A., Fabeni,L., Butera,O., Colavita,F., Meschi,S., Matusali,G., Rivano Capparuccia,M., Specchiarello,E., Vairo,F., Vaia,F., Nicastri,E., Antinori,A., Girardi,E. and Maggi,F. |
| EPI_ISL_13331714 | Department of Virology, Faculty of Medicine, University of Helsinki, Hartmaninkatu 3 | Department of Virology, Faculty of Medicine, University of Helsinki, Hartmaninkatu 3 | Kant,R., Smura,T., Vauhkonen,H. and Vapalahti,O. |
| EPI_ISL_13331715 | Department of Virology, Faculty of Medicine, University of Helsinki, Hartmaninkatu 3 | Department of Virology, Faculty of Medicine, University of Helsinki, Hartmaninkatu 3 | Kant,R., Smura,T., Vauhkonen,H., Vapalahti,O. and Sironen,T. |
| EPI_ISL_13331717 | Genomics Division, Instituto Tecnologico y de Energias Renovables (ITER), Poligono Industrial de Granadilla | Genomics Division, Instituto Tecnologico y de Energias Renovables (ITER), Poligono Industrial de Granadilla, | Alcoba-Florez,J., Munoz-Barrera,A., Ciuffreda,L., Rodriguez-Perez,H., Rubio-Rodriguez,L.A., Gil-Campesino,H., Garcia-Martinez de Artola,D., Inigo-Campos,A., Diez-Gil,O., Gonzalez-Montelongo,R., Valenzuela-Fernandez,A., Lorenzo-Salazar,J.M. and Flores,C. |
| EPI_ISL_13338028 | Clinical Virology Unit, Department of Clinical Sciences, Institute of Tropical Medicine of Antwerp | Clinical Virology Unit, Department of Clinical Sciences, Institute of Tropical Medicine of Antwerp | Antonio Mauro Rezende*, Tessa de Block*, Sandra Coppens, Eric Florence, Maartje van Frankenhuijsen, Stefanie Bracke, Isabel Brosius, Laurens Liesenborghs, Patrick Soentjens, Kevin Ariën, Marjan Van Esbroeck, Philippe Selhorst*, Koen Vercauteren* *equal contribution |
| EPI_ISL_13339105 | Microbiology Service, Hospital Universitario Clinico San Cecilio, Granada | Microbiology Service, Hospital Universitario Clinico San Cecilio, Granada | Chueca N, de Salazar A, Viñuela L, Fuentes A, Casimiro-Soriguer CS, Perez-Florido J, Dopazo J, Garcia F |
| EPI_ISL_13342823 | Clinical Virology Unit, Department of Clinical Sciences, Institute of Tropical Medicine of Antwerp | Clinical Virology Unit, Department of Clinical Sciences, Institute of Tropical Medicine of Antwerp | Philippe Selhorst, Antonio Mauro Rezende, Tessa de Block, Sandra Coppens, Eric Florence, Isabel Brosius, Laurens Liesenborghs, Kevin Ariën, Marjan Van Esbroeck, Chris Kenyon, Koen Vercauteren |
| EPI_ISL_13343634 | Instituto de Infectologia Emilio Ribas | Instituto Adolfo Lutz Strategic Laboratory | Claudio Tavares Sacchi, Karoline Rodrigues Campos, Adriano Abbud, Adriana Bugno |
| EPI_ISL_13343697 | Fleury Medicina Dignóstica | Instituto Adolfo Lutz Strategic Laboratory | Claudio Tavares Sacchi, Karoline Rodrigues Campos, Adriano Abbud, Adriana Bugno |
| EPI_ISL_13343718 | Hospital Santa Ignes | Instituto Adolfo Lutz Strategic Laboratory | Claudio Tavares Sacchi, Karoline Rodrigues Campos, Adriano Abbud, Adriana Bugno |
| EPI_ISL_13351002 | B.C. Centre for Disease Control Public Health Laboratory | B.C. Centre for Disease Control Public Health Laboratory | John Tyson, Tracy Lee, Anthea Lam, Josh Quick, Agatha Jassem, Natalie Prystajecky, Linda Hoang, Inna Sekirov, Catherine Hogn, Frankie Tsang, Mel Krajden |
| EPI_ISL_13362760, EPI_ISL_13362764 | Laboratorio di Epidemiologia Molecolare e Sanità Pubblica-Policlinico Bari | Istituto Zooprofilattico Sperimentale della Puglia e della Basilicata | Parisi A, Simone D, Capozzi L, Del Sambro L, Bianco A, Chironna M, Loconsole D, Sallustio F, Galante D, Pace L, Manzulli V, Fasanella A. |
| EPI_ISL_13363142 | Hospital Universitari Vall d'Hebron | Hospital Universitari Vall d'Hebron | Maria Piñana, Cristina Andrés, Alejandra González-Sánchez, Damir Garcia-Cehic, Ariadna Rando, Juliana Esperalba, Maria Gema Codina, Maria Carmen Martin, Carla Castillo, Karen García, Rodrigo Vásquez, Maria Piquer, Tomàs Pumarola, Josep Quer, Andrés Antón |
| EPI_ISL_13374487 | National Public Health Center, National Biosafety Laboratory | National Public Health Center, National Biosafety Laboratory | Judit Henczkó, Dániel Déri, Fruzsina Petrovay, Lili Jármi, Bernadett Pályi, Eszter Balla, Zoltán Kis |
| EPI_ISL_13408799, EPI_ISL_13408801, EPI_ISL_13408803 | Public Health Agency of Canada, National Microbiology Laboratory | Public Health Agency of Canada, National Microbiology Laboratory | Knox,N., Hole,D., Duggan,A., Yadav,C., Haidl,E., Chapel,M., Graham,M., Domselaar,G.V., Jolly,G., Audet,J., Fernando,L., Antonation,K., Hagan,M., Griffiths,E., Leung,A., Safronetz,D., Eshaghi,A., Gubbay,J.B., Hasso,M., Marchand-Austin,A., Olsha,R. and Patel,S.N. |
| EPI_ISL_13408805, EPI_ISL_13408807, EPI_ISL_13408809, EPI_ISL_13408811, EPI_ISL_13408813, EPI_ISL_13408815, EPI_ISL_13408817, EPI_ISL_13408819, EPI_ISL_13408821, EPI_ISL_13408823, EPI_ISL_13408825, EPI_ISL_13408827, EPI_ISL_13408829, EPI_ISL_13408831, EPI_ISL_13408833, EPI_ISL_13408835 | | | |
| see above | Public Health Agency of Canada, National Microbiology Laboratory | Public Health Agency of Canada, National Microbiology Laboratory | ncknox |
| EPI_ISL_13408837, EPI_ISL_13408839, EPI_ISL_13408841, EPI_ISL_13408843, EPI_ISL_13408845, EPI_ISL_13408847, EPI_ISL_13408849, EPI_ISL_13408851, EPI_ISL_13408853, EPI_ISL_13408855, EPI_ISL_13408857, EPI_ISL_13408859, EPI_ISL_13408861 | | | |
| see above | Public Health Agency of Canada, National Microbiology Laboratory | Public Health Agency of Canada, National Microbiology Laboratory | Knox,N., Duggan,A., Yadav,C., Hole,D., Haidl,E., Chapel,M., Jolly,G., Domselaar,G.V., Antonation,K., Leung,A., Fernando,L., Audet,J., Hagan,M., Graham,M., Griffiths,E., Safronetz,D., Charest,H., Levade,I. and Fafard,J. |
| EPI_ISL_13409177, EPI_ISL_13409178, EPI_ISL_13409179, EPI_ISL_13409180, EPI_ISL_13409181 | Viral Genomics and Bioinformatics, MRC University of Glasgow Centre for Virus Research | Viral Genomics and Bioinformatics, MRC University of Glasgow Centre for Virus Research | Filipe,A., Tong,L., Vattipally,S.B., Maclean,A., Gunson,R., Holden,M.T.G., Barr,D., Ho,A., Palmarini,M., Rambaut,A., Robertson,D.L. and Thomson,E.C. |
| EPI_ISL_13411153, EPI_ISL_13411154, EPI_ISL_13411155, EPI_ISL_13411156, EPI_ISL_13411157, EPI_ISL_13411158 | Centre for Biological Threats, Highly Pathogenic Viruses, Robert Koch Institute | Centre for Biological Threats, Highly Pathogenic Viruses, Robert Koch Institute | Brinkmann,A., Kohl,C., Uddin,S., Pape,K., Schrick,L., Michel,J., Schaade,L. and Nitsche,A. |
| EPI_ISL_13411159, EPI_ISL_13411160, EPI_ISL_13411161, EPI_ISL_13411162 | Centre for Biological Threats, Highly Pathogenic Viruses, Robert Koch Institute | Centre for Biological Threats, Highly Pathogenic Viruses, Robert Koch Institute | Brinkmann,A., Kohl,C., Uddin,S., Pape,K., Schrick,L., Michel,J., Stocker,H., Schaade,L. and Nitsche,A. |
| EPI_ISL_13411163, EPI_ISL_13411164, EPI_ISL_13411165 | Centre for Biological Threats, Highly Pathogenic Viruses, Robert Koch Institute | Centre for Biological Threats, Highly Pathogenic Viruses, Robert Koch Institute | Brinkmann,A., Kohl,C., Uddin,S., Pape,K., Schrick,L., Michel,J., Schaade,L. and Nitsche,A. |
| EPI_ISL_13411166, EPI_ISL_13411167 | Centre for Biological Threats, Highly Pathogenic Viruses, Robert Koch Institute | Centre for Biological Threats, Highly Pathogenic Viruses, Robert Koch Institute | Brinkmann,A., Kohl,C., Uddin,S., Pape,K., Schrick,L., Michel,J., Jessen,H., Schaade,L. and Nitsche,A. |
| EPI_ISL_13411168 | Centre for Biological Threats, Highly Pathogenic Viruses, Robert Koch Institute | Centre for Biological Threats, Highly Pathogenic Viruses, Robert Koch Institute | Brinkmann,A., Kohl,C., Uddin,S., Pape,K., Schrick,L., Michel,J., Pfaefflin,F., Schaade,L. and Nitsche,A. |
| EPI_ISL_13436658 | Coordenadoria de Vigilancia em Saude - Sao Paulo | Instituto Adolfo Lutz Strategic Laboratory | Claudio Tavares Sacchi, Karoline Rodrigues Campos, Ariadne Fereira Amarante, Adriano Abbud, Adriana Bugno |
| EPI_ISL_13436792 | Hospital Santa Ignes | Instituto Adolfo Lutz Strategic Laboratory | Claudio Tavares Sacchi, Karoline Rodrigues Campos, Adriano Abbud, Adriana Bugno |
| EPI_ISL_13437056 | Hosp. Alemao Oswaldo Cruz | Instituto Adolfo Lutz Strategic Laboratory | Claudio Tavares Sacchi, Karoline Rodrigues Campos, Ariadne Ferreira Amarante, Adriano Abbud, Adriana Bugno |

| | | | |
|---|---|---|---|
| EPI_ISL_13445553 | Laboratory for Diagnostics of Zoonoses and WHO Centre, Institute of Microbiology and Immunology, Faculty of Medicine, University of Ljubljana | Laboratory for Diagnostics of Zoonoses and WHO Centre, Institute of Microbiology and Immunology, Faculty of Medicine, University of Ljubljana | Zakotnik,S., Vlaj,D., Suljic,A., Zorec,T.M., Korva,M., Poljak,M. and Avsic Zupanc,T. |
| EPI_ISL_13449965, EPI_ISL_13449966 | Hospital Universitario La Paz, Microbiology | Hospital Universitario La Paz, Microbiology | de la Hoz-Sanchez,B., Lopez-Ortiz,M., Gutierrez-Arroyo,A., Roces-Alvarez,P., Lazaro-Peona,F., Dahdouh,E., Bloise,I., Garcia-Rodriguez,J. and Mingorance,J. |
| EPI_ISL_13459346 | CRT-DST-AIDS | Instituto Adolfo Lutz Strategic Laboratory | Claudio Tavares Sacchi, Karoline Rodrigues Campos, Ariadne Ferreira Amarante, Adriano Abbud, Adriana Bugno |
| EPI_ISL_13459347, EPI_ISL_13459482, EPI_ISL_13459483 | Instituto de Infectologia Emilio Ribas | Instituto Adolfo Lutz Strategic Laboratory | Claudio Tavares Sacchi, Karoline Rodrigues Campos, Ariadne Ferreira Amarante, Adriano Abbud, Adriana Bugno |
| EPI_ISL_13466446, EPI_ISL_13466447, EPI_ISL_13466448, EPI_ISL_13466449, EPI_ISL_13466450, EPI_ISL_13466451, EPI_ISL_13466452, EPI_ISL_13466453, EPI_ISL_13466454, EPI_ISL_13466455, EPI_ISL_13466456, EPI_ISL_13466457, EPI_ISL_13466458, EPI_ISL_13466459, EPI_ISL_13466460, EPI_ISL_13466461, EPI_ISL_13466462, EPI_ISL_13466463, EPI_ISL_13466464, EPI_ISL_13466465 | | | |
| see above | Department of Infectious Diseases, National Institute of Health Doutor Ricardo Jorge, Portugal (INSA) | Department of Infectious Diseases, National Institute of Health Doutor Ricardo Jorge, Portugal (INSA) | Isidro,J., Borges,V., Pinto,M., Sobral,D., Santos,J., Nunes,A., Mixao,V., Ferreira,R., Santos,D., Duarte,S., Vieira,L., Borrego,M.J., Nuncio,S., Lopes de Carvalho,I., Pelerito,A., Cordeiro,R., Gomes,J.P. |
| EPI_ISL_13472080 | National Institute of Public Health NIH - NRI | National Institute of Public Health NIH - NRI | Wokowicz Tomasz, Zacharczuk Katarzyna, Gierczyski Rafa |
| EPI_ISL_13472250 | Medical University of Vienna Center for Virology | Medical University of Vienna Center for Virology | Jeremy V. Camp, Monika Redlberger-Fritz, Stephan W. Aberle |
| EPI_ISL_13483155, EPI_ISL_13483157, EPI_ISL_13483159, EPI_ISL_13483161, EPI_ISL_13483162, EPI_ISL_13483163, EPI_ISL_13483164 | Centre for Biological Threats, Highly Pathogenic Viruses, Robert Koch Institute | Centre for Biological Threats, Highly Pathogenic Viruses, Robert Koch Institute | Brinkmann,A., Kohl,C., Uddin,S., Pape,K., Schrick,L., Michel,J., Jessen,H., Schaade,L. and Nitsche,A. |
| EPI_ISL_13483165, EPI_ISL_13483167, EPI_ISL_13483168, EPI_ISL_13483170, EPI_ISL_13483171, EPI_ISL_13483173, EPI_ISL_13483175, EPI_ISL_13483177, EPI_ISL_13483178, EPI_ISL_13483180, EPI_ISL_13483182, EPI_ISL_13483183, EPI_ISL_13483185, EPI_ISL_13483187, EPI_ISL_13483188, EPI_ISL_13483190, EPI_ISL_13483191, EPI_ISL_13483193, EPI_ISL_13483195, EPI_ISL_13483196, EPI_ISL_13483198, EPI_ISL_13483200, EPI_ISL_13483201, EPI_ISL_13483203, EPI_ISL_13483205, EPI_ISL_13483206, EPI_ISL_13483208 | | | |
| see above | Centre for Biological Threats, Highly Pathogenic Viruses, Robert Koch Institute | Centre for Biological Threats, Highly Pathogenic Viruses, Robert Koch Institute | Brinkmann,A., Kohl,C., Uddin,S., Pape,K., Schrick,L., Michel,J., Schaade,L. and Nitsche,A. |
| EPI_ISL_13483780 | UCD National Virus Reference Laboratory, University College Dublin | UCD National Virus Reference Laboratory, University College Dublin | Nicola Fletcher, Gabriel Gonzalez, Luke Meredith, Kevin Purves, Michael Carr, Jonathan Dean, Brian Keogan, Brendan Crowley, Fiona Lyons, Sophie O'Reilly, Virginie Gautier, Patrick Mallon, Stephen Gordon, Jeff Connell, Cillian F De Gascun |
| EPI_ISL_13484458 | Laboratorio de Enterovirus, Instituto Oswaldo Cruz, Fiocruz | Instituto Oswaldo Cruz FIOCRUZ - Laboratory of Respiratory Viruses and Measles (LVRS) | Paola Resende, Elisa Cavalcante Pereira, Bruna Mendonça da Silva, Jéssica Graça Macedo de Carvalho, Larissa Macedo Pinto, Victor Guimaraes, Marilda Siqueira, Renan da Silva Faustino, Marilia Santini, Edson Elias da Silva on behalf of the Fiocruz Genomic Surveillance Network |
| EPI_ISL_13498265 | National Institute for Communicable Diseases of the National Health Laboratory Service | National Institute for Communicable Diseases of the National Health Laboratory Service | Chan WY, Mtshali PS, Grobbelaar A, Moolla N, Mohale T, Du Plessis MG, Ismail A, Weyer J |
| EPI_ISL_13499566 | Erasmus Medical Center Department of Virology | Erasmus Medical Center Department of Virology | Bas Oude Munnink, Marjan Boter, Babette Weller, Richard Molenkamp, Janette Rahamat-Langendoen, Reina Sikkema, Marion Koopmans |
| EPI_ISL_13502582 | Laboratory of Microbiology and Virology, Ospedale Amedeo di Savoia, ASL "Città di Torino" | Laboratory of Microbiology and Virology, Ospedale Amedeo di Savoia, ASL "Città di Torino" | Francesco Cerutti, Antonella Bottoni, Marisa Cazzadore, Tiziano Allice, Maria Grazia Milia, Gabriella Gregori, Elisa Burdino, Valeria Ghisetti |
| EPI_ISL_13508393 | Hosp. Itacolomy Butanta | Instituto Adolfo Lutz Strategic Laboratory | Claudio Tavares Sacchi, Karoline Rodrigues Campos, Ariadne Ferreira Amarante, Adriano Abbud, Adriana Bugno |
| EPI_ISL_13508471 | Instituto de Infectologia Emilio Ribas | Instituto Adolfo Lutz Strategic Laboratory | Claudio Tavares Sacchi, Karoline Rodrigues Campos, Ariadne Ferreira Amarante, Adriano Abbud, Adriana Bugno |
| EPI_ISL_13511312 | Laboratorio de Salud Pública de Antioquia | Instituto Nacional de Salud- Dirección de Investigación en Salud Pública | Katherine Laiton-Donato, Diego A. Álvarez-Díaz, Carlos Franco-Muñoz, Héctor A. Ruiz-Moreno, Paola Rojas-Estevez, Andres Prada, Alicia Rosales, Marcela Mercado-Reyes |
| EPI_ISL_13530881 | Laboratorio de Referencia Nacional de Virus Respiratorios. Centro Nacional de Salud Publica. Instituto Nacional de Salud Peru. | Laboratorio de Referencia Nacional de Virus Respiratorios. Centro Nacional de Salud Publica. Instituto Nacional de Salud Peru. | Carlos Padilla Rojas, Veronica Hurtado Vela, Iris Silva Molina, Luren Sevilla Castañeda, Victor Jimenez Vasquez, Orson Mestanza Millones, Luis Barcena Flores, Wendy Lizarraga Olivares, Alicia Nuñez Llanos, Steve Acedo Lazo, Francisco Ascue Orosco, Kelly Izarra Rojas, Princesa Medrano Alhuay, Karla Vasquez Cajachahua, Estela Huaman Angeles, Jorge Giraldo Chavez, Lilian Huarca Balbin, Lisbet Roxana Inga Angulo, Maria Sandra Villar Saavedra, Henri Bailon Calderon, Lely Solari Zerpa, Gloria Arotinco Garayar. Equipo de vigilancia genomica del Instituto Nacional de Salud. |
| EPI_ISL_13537922 | Instituto de Medicina Tropical de Sao Paulo (IMT-USP) | School of Public Health, Imperial College London | Coletti,T.M., Ghilardi,F., khan,M.J., Claro,I.M., Valenca,I.N., Faria,N.R. and Sabino,E.C. |
| EPI_ISL_13537923 | Microbiology, Immunology and Transplantation, KU Leuven, Rega Institute | Microbiology, Immunology and Transplantation, KU Leuven, Rega Institute | Wawina-Bokalanga,T., Vanmechelen,B., Logist,A.-S., Sinnesael,R., Ysebaert,L., Bloemen,M. and Maes,P. |
| EPI_ISL_13537924, EPI_ISL_13537925, EPI_ISL_13537926 | Microbiology, Immunology and Transplantation, KU Leuven, Rega Institute | Microbiology, Immunology and Transplantation, KU Leuven, Rega Institute | Vanmechelen,B., Wawina-Bokalanga,T., Logist,A.-S., Sinnesael,R., Ysebaert,L., Verlinden,J., Van Holm,B., Bloemen,M. and Maes,P. |
| EPI_ISL_13544223, EPI_ISL_13544224, EPI_ISL_13544225, EPI_ISL_13544226, EPI_ISL_13544227, EPI_ISL_13544228, EPI_ISL_13544229, EPI_ISL_13544230, EPI_ISL_13544231, EPI_ISL_13544232, EPI_ISL_13544233, EPI_ISL_13544234, EPI_ISL_13544235, EPI_ISL_13544236 | | | |
| see above | Public Health Agency of Canada, National Microbiology Laboratory | Public Health Agency of Canada, National Microbiology Laboratory | Duggan,A., Hole,D., Knox,N., Yadav,C., Haidl,E., Chapel,M., Domselaar,G.V., Jolly,G., Audet,J., Fernando,L., Antonation,K., Safronetz,D., Hagan,M., Griffiths,E., Leung,A., Graham,M., Peters,G., Go,A., Laminman,V., Kaplen,B., Eshaghi,A., Gubbay,J.B., Hasso,M., Marchand-Austin,A., Olsha,R. and Patel,S.N. |
| EPI_ISL_13544237, EPI_ISL_13544238, EPI_ISL_13544239, EPI_ISL_13544240, EPI_ISL_13544241, EPI_ISL_13544242, EPI_ISL_13544243, EPI_ISL_13544244, EPI_ISL_13544245, EPI_ISL_13544246, EPI_ISL_13544247, EPI_ISL_13544248, EPI_ISL_13544249, EPI_ISL_13544250, EPI_ISL_13544251, EPI_ISL_13544252, EPI_ISL_13544253, EPI_ISL_13544254, EPI_ISL_13544255, EPI_ISL_13544256, EPI_ISL_13544257, EPI_ISL_13544258, EPI_ISL_13544259, EPI_ISL_13544260, EPI_ISL_13544261, EPI_ISL_13544262, EPI_ISL_13544263, EPI_ISL_13544264, EPI_ISL_13544265, EPI_ISL_13544266, EPI_ISL_13544267 | | | |
| see above | Public Health Agency of Canada, National Microbiology Laboratory | Public Health Agency of Canada, National Microbiology Laboratory | Duggan,A., Hole,D., Knox,N., Yadav,C., Haidl,E., Chapel,M., Domselaar,G.V., Fernando,L., Graham,M., Antonation,K., Audet,J., Hagan,M., Safronetz,D., Leung,A., Peters,G., Go,A., Laminman,V., Kaplen,B., Jolly,G., Charest,H., Levade,I. and Fafard,J. |
| EPI_ISL_13573943 | Center for Virology, Medical University of Vienna | Medical University of Vienna Center for Virology | Jeremy V. Camp, Monika Redlberger-Fritz, Stephan W. Aberle |
| EPI_ISL_13584854, EPI_ISL_13586184 | Institute for Virology, Philipps-University Marburg | Institute for Virology, Philipps-University Marburg | Eickmann, M., Lier, C., Kowalski, K., Kraft, F., Becker, S. |
| EPI_ISL_13624509 | Instituto de Diagnóstico y Referencia Epidemiológicos/Jurisdicción Sanitaria Cuauhtémoc/Hospital Ángeles Roma | Instituto de Diagnóstico y Referencia Epidemiológicos/Instituto de Biotecnología UNAM | Adnan Araiza-Rodríguez, Adriana Salvador-Patiño, Alejandro Sánchez-Flores, América del Pilar Mandujano-Martínez, Blanca Taboada, Carlos Eduardo Hernández-Sánchez, Carlos F. Arias, Claudia Elena Wong-Arámbula, Daniel José Regalado-Santiago, David Esaú Fragoso-Fonseca, Elizabeth Andrade-Montiel, Fabiola Garcés-Ayala, Fernando González-Dominguez, Gabriel García-Rodríguez, Gloria Vázquez-Castro, Hugo López Gatell Ramírez, Irma López-Martínez, Jerome Verleyen, Jesús Trujillo, Jorge Ochoa, José Ernesto Ramírez-González, Karel Estrada-Guerra, Lucía Hernández-Rivas, Magaly Guadalupe Landa-Flores, Maribel González-Villa, Mireya Mederos-Michel, Nancy Martínez-Velázquez, Noé Escobar-Escamilla, Oliva López, Ricardo Cortés-Alcalá, Ricardo Grande, Verónica Jiménez-Jacinto |
| EPI_ISL_13632068 | SA Pathology | SA Pathology | Coldbeck-Shackley, R, Selway, C, Adamson, PJ, Lim, CK, Turra, M, Bastian, I, Beazley, R, Flood, L, Leong, LEX |
| EPI_ISL_13632071 | Center of Diagnostics and Vaccine Development, Centers for Disease Control, Taiwan | Center of Diagnostics and Vaccine Development, Centers for Disease Control, Taiwan | Jih-Hui Lin, Shu-Chun Chiu, Hsin-I, Huang, Wei-Lun Huang, Wen-Bin, Fann, Pei-Yu, Hsieh, Jyh-Yuan Yang |
| EPI_ISL_13632288 | National Institute for Communicable Diseases of the National Health Laboratory Service | National Institute for Communicable Diseases of the National Health Laboratory Service | Chan WY, Mtshali PS, Grobbelaar A, Moolla N, Mohale T, Lowe M, Du Plessis MG, Ismail A, Weyer J |
| EPI_ISL_13651348, EPI_ISL_13651349, EPI_ISL_13651350 | Laboratorio de Referencia Nacional de Virus Respiratorio. Centro Nacional de Salud Publica. Instituto Nacional de Salud. | Laboratorio de Referencia Nacional de Virus Respiratorio. Centro Nacional de Salud Publica. Instituto Nacional de Salud. | Carlos Padilla Rojas, Veronica Hurtado Vela, Iris Silva Molina, Luren Sevilla Castañeda, Victor Jimenez Vasquez, Orson Mestanza Millones, Luis Barcena Flores, Wendy Lizarraga Olivares, Alicia Nuñez Llanos, Steve Acedo Lazo, Francisco Ascue Orosco, Kelly Izarra Rojas, Princesa Medrano Alhuay, Karla Vasquez Cajachahua, Estela Huaman Angeles, Jorge Giraldo Chavez, Lilian Huarca Balbin, Lisbet Roxana Inga Angulo, Maria Sandra Villar Saavedra, Henri Bailon Calderon, Lely Solari Zerpa, Gloria Arotinco Garayar. Equipo de vigilancia genomica del Instituto Nacional de Salud. |
| EPI_ISL_13658019, EPI_ISL_13658021 | Erasmus Medical Center Department of Virology | Erasmus Medical Center Department of Virology | Bas Oude Munnink, Marjan Boter, Babette Weller, Richard Molenkamp, Janette Rahamat-Langendoen, Reina Sikkema, Marion Koopmans |

| | | | |
|---|---|---|---|
| EPI_ISL_13660191 | Hospital Center Luxembourg | Laboratoire National de Santé Microbiology | Eric Hugoson, Ines Kozar, Sibel Berger, Anke Wienecke-Baldacchino, Bas Oude Munnink, Michel Kohnen, Jean-Hugues Francois, Tamir Abdelrahman |
| EPI_ISL_13705358 | Hosp. Alemao Oswaldo Cruz | Instituto Adolfo Lutz Strategic Laboratory | Claudio Tavares Sacchi, Karoline Rodrigues Campos, Ariadne Ferreira Amarante, Marlon Benedito Nascimento Santos, Alex Domingos Reis, Adriano Abbud, Adriana Bugno |
| EPI_ISL_13705407 | Hosp. Sirio-Libanes | Instituto Adolfo Lutz Strategic Laboratory | Claudio Tavares Sacchi, Karoline Rodrigues Campos, Ariadne Ferreira Amarante, Marlon Benedito Nascimento Santos, Alex Domingos Reis, Adriano Abbud, Adriana Bugno |
| EPI_ISL_13717674 | Hospital Center Luxembourg | Laboratoire National de Santé Microbiology | Eric Hugoson, Ines Kozar, Sibel Berger, Anke Wienecke-Baldacchino, Bas Oude Munnink, Michel Kohnen, Jean-Hugues Francois, Tamir Abdelrahman |
| EPI_ISL_13728303 | Department of Medical Microbiology & Infection prevention, Amsterdam University Medical Centers location AMC | Department of Medical Microbiology & Infection prevention, Amsterdam University Medical Centers location AMC | Matthijs Welkers, Jelle Koopsen, Robin van Houdt, Marcel Jonges, Sebastien Matamoros, Sjoerd Rebers, Fokla Zorgdrager, Sylvia Bruisten, Judith den Uil, Akke Cornelissen, Janke Schinkel, Menno de Jong, Gini van Rijckevorsel and Mariken van der Lubben on behalf of the Amsterdam Regional Genomic epidemiology and Outbreak Surveillance (ARGOS) consortium |
| EPI_ISL_13732932 | Hosp. Sao Joaquim - Beneficiencia Portuguesa | Instituto Adolfo Lutz Strategic Laboratory | Claudio Tavares Sacchi, Karoline Rodrigues Campos, Ariadne Ferreira Amarante, Marlon Benedito Nascimento Santos, Alex Domingos Reis, Adriano Abbud, Adriana Bugno |
| EPI_ISL_13734233 | Microbial Genomics, Hospital General Universitario Gregorio Maranon | Microbial Genomics, Hospital General Universitario Gregorio Maranon | Palomino-Cabrera,R., Penas-Utrilla,D., Buenestado-Serrano,S., Perez-Lago,L., Herranz Martin,M., Veintimilla,C., Catalan,P., Munoz,P. and Garcia de Viedma,D. |
| EPI_ISL_13734237, EPI_ISL_13734238, EPI_ISL_13734239, EPI_ISL_13734240, EPI_ISL_13734241, EPI_ISL_13734242, EPI_ISL_13734243, EPI_ISL_13734244, EPI_ISL_13734245, EPI_ISL_13734246, EPI_ISL_13734247, EPI_ISL_13734248, EPI_ISL_13734249, EPI_ISL_13734250, EPI_ISL_13734251, EPI_ISL_13734252, EPI_ISL_13734253, EPI_ISL_13734254, EPI_ISL_13734255, EPI_ISL_13734256, EPI_ISL_13734257, EPI_ISL_13734258, EPI_ISL_13734259, EPI_ISL_13734260, EPI_ISL_13734261, EPI_ISL_13734262, EPI_ISL_13734263, EPI_ISL_13734264, EPI_ISL_13734265, EPI_ISL_13734266, EPI_ISL_13734267, EPI_ISL_13734268 | | | |
| see above | Centre for Biological Threats, Highly Pathogenic Viruses, Robert Koch Institute | Centre for Biological Threats, Highly Pathogenic Viruses, Robert Koch Institute | Brinkmann,A., Kohl,C., Pape,K., Uddin,S., Schrick,L., Michel,J., Schaade,L. and Nitsche,A. |
| EPI_ISL_13734269 | Department of Clinical Sciences, Institute of Tropica Medicine | Department of Clinical Sciences, Institute of Tropica Medicine | De Baetselier,I., Van Dijck,C., Kenyon,C., Coppens,J., Smet,H., de Block,T., Coppens,S., Vanroye,F., Bugert,J., Girl,P., Liesenborghs,L., Selhorst,P., Arien,K., Van den Bossche,D., Florence,E., Rezende,A.M., Vercauteren,K. and Van Esbroeck,M. |
| EPI_ISL_13734270 | Centers for Disease Control & Prevention (CDC), Division of High Consequence Pathogens and Pathology (DHCPP-PRB) | Centers for Disease Control & Prevention (CDC), Division of High Consequence Pathogens and Pathology (DHCPP-PRB) | Gigante,C.M., Ventura,J., Seabolt,M.H., Zhao,H., Wilkins,K., Respress,J., Howard,D., Batra,D., McCollum,A., Hutson,C., Davidson,W., Rao,A., Nash,J. and Li,Y. |
| EPI_ISL_13744896 | Centers for Disease Control & Prevention (CDC), Division of High Consequence Pathogens and Pathology (DHCPP-PRB) | Centers for Disease Control & Prevention (CDC), Division of High Consequence Pathogens and Pathology (DHCPP-PRB) | Gigante,C.M., Ghinai,I., Seabolt,M.H., Zhao,H., Wilkins,K., Respress,J., Howard,D., Batra,D., McCollum,A., Hutson,C., Davidson,W., Rao,A., Kerins,J. and Li,Y. |
| EPI_ISL_13744897, EPI_ISL_13744898 | Centers for Disease Control & Prevention (CDC), Division of High Consequence Pathogens and Pathology (DHCPP-PRB) | Centers for Disease Control & Prevention (CDC), Division of High Consequence Pathogens and Pathology (DHCPP-PRB) | Gigante,C.M., Hughes,S., Seabolt,M.H., Zhao,H., Wilkins,K., Respress,J., Howard,D., Batra,D., McCollum,A., Hutson,C., Davidson,W., Rao,A., Baumgartner,J. and Li,Y. |
| EPI_ISL_13744899 | Centers for Disease Control & Prevention (CDC), Division of High Consequence Pathogens and Pathology (DHCPP-PRB) | Centers for Disease Control & Prevention (CDC), Division of High Consequence Pathogens and Pathology (DHCPP-PRB) | Gigante,C.M., Ghinai,I., Seabolt,M.H., Zhao,H., Wilkins,K., Respress,J., Howard,D., Batra,D., McCollum,A., Hutson,C., Davidson,W., Rao,A., Kerins,J. and Li,Y. |
| EPI_ISL_13744900, EPI_ISL_13744901 | Centers for Disease Control & Prevention (CDC), Division of High Consequence Pathogens and Pathology (DHCPP-PRB) | Centers for Disease Control & Prevention (CDC), Division of High Consequence Pathogens and Pathology (DHCPP-PRB) | Gigante,C.M., Hughes,S., Seabolt,M.H., Zhao,H., Wilkins,K., Respress,J., Howard,D., Batra,D., McCollum,A., Hutson,C., Davidson,W., Rao,A., Baumgartner,J. and Li,Y. |
| EPI_ISL_13744902 | Department of Virology, Faculty of Medicine, University of Helsinki | Department of Virology, Faculty of Medicine, University of Helsinki | Kant,R., Smura,T., Vauhkonen,H. and Vapalahti,O. |
| EPI_ISL_13744903, EPI_ISL_13744904, EPI_ISL_13744905 | Centre for Biological Threats, Highly Pathogenic Viruses, Robert Koch Institute | Centre for Biological Threats, Highly Pathogenic Viruses, Robert Koch Institute | Brinkmann,A., Kohl,C., Pape,K., Uddin,S., Schrick,L., Michel,J., Jessen,H., Schaade,L. and Nitsche,A. |
| EPI_ISL_13744906, EPI_ISL_13744907, EPI_ISL_13744908, EPI_ISL_13744909, EPI_ISL_13744910, EPI_ISL_13744911, EPI_ISL_13744912, EPI_ISL_13744913, EPI_ISL_13744914, EPI_ISL_13744915, EPI_ISL_13744916, EPI_ISL_13744917, EPI_ISL_13744918, EPI_ISL_13744919, EPI_ISL_13744920, EPI_ISL_13744921, EPI_ISL_13744922, EPI_ISL_13744923, EPI_ISL_13744924, EPI_ISL_13744925, EPI_ISL_13744926, EPI_ISL_13744927, EPI_ISL_13744928, EPI_ISL_13744929, EPI_ISL_13744930, EPI_ISL_13744931 | | | |
| see above | Centre for Biological Threats, Highly Pathogenic Viruses, Robert Koch Institute | Centre for Biological Threats, Highly Pathogenic Viruses, Robert Koch Institute | Brinkmann,A., Kohl,C., Pape,K., Uddin,S., Schrick,L., Michel,J., Schaade,L. and Nitsche,A. |
| EPI_ISL_13822667, EPI_ISL_13822668, EPI_ISL_13822669, EPI_ISL_13822718 | Erasmus Medical Center Department of Virology | Erasmus Medical Center Department of Virology | Bas Oude Munnink, Marjan Boter, Babette Weller, Richard Molenkamp, Janette Rahamat-Langendoen, Reina Sikkema, Marion Koopmans |
| EPI_ISL_13827273, EPI_ISL_13827274, EPI_ISL_13827275, EPI_ISL_13827277, EPI_ISL_13827278, EPI_ISL_13827279, EPI_ISL_13827280, EPI_ISL_13827281, EPI_ISL_13827282 | Public Health Agency of Canada, National Microbiology Laboratory | Public Health Agency of Canada, National Microbiology Laboratory | Duggan,A., Hole,D., Yadav,C., Knox,N., Haidl,E., Chapel,M., Domselaar,G.V., Fernando,L., Graham,M., Antonation,K., Audet,J., Hagan,M., Safronetz,D., Leung,A., Peters,G., Go,A., Laminman,V., Kaplen,B., Jolly,G., Marchand-Austin,A., Eshaghi,A., Patel,S.N., Hasso,M., Gubbay,J.B. and Olsha,R. |
| EPI_ISL_13833194, EPI_ISL_13833195, EPI_ISL_13833196, EPI_ISL_13833197 | Laboratorio de Referencia Nacional de Virus Respiratorio. Centro Nacional de Salud Publica. Instituto Nacional de Salud. | Laboratorio de Referencia Nacional de Virus Respiratorio. Centro Nacional de Salud Publica. Instituto Nacional de Salud. | Carlos Padilla Rojas, Veronica Hurtado Vela, Iris Silva Molina, Luren Sevilla Castañeda, Victor Jimenez Vasquez, Orson Mestanza Millones, Luis Barcena Flores, Wendy Lizarraga Olivares, Alicia Nuñez Llanos, Steve Acedo Lazo, Francisco Ascue Orosco, Kelly Izarra Rojas, Princesa Medrano Alhuay, Karla Vasquez Cajachahua, Estela Huaman Angeles, Jorge Giraldo Chavez, Lilian Huarca Balbin, Lisbet Roxana Inga Angulo, Maria Sandra Villar Saavedra, Henri Bailon Calderon, Lely Solari Zerpa, Gloria Arotinco Garayar. Equipo de vigilancia genomica del Instituto Nacional de Salud. |
| EPI_ISL_13842269, EPI_ISL_13842548 | Center for Virology, Medical University of Vienna | Medical University of Vienna Center for Virology | Jeremy V. Camp, Monika Redlberger-Fritz, Stephan W. Aberle |
| EPI_ISL_13889435, EPI_ISL_13889436, EPI_ISL_13889437, EPI_ISL_13889438, EPI_ISL_13889439, EPI_ISL_13889440, EPI_ISL_13889441, EPI_ISL_13889442, EPI_ISL_13889443, EPI_ISL_13889444, EPI_ISL_13889445, EPI_ISL_13889446, EPI_ISL_13889447, EPI_ISL_13889448, EPI_ISL_13889449, EPI_ISL_13889450, EPI_ISL_13889515, EPI_ISL_13889590, EPI_ISL_13889660, EPI_ISL_13889729, EPI_ISL_13889796, EPI_ISL_13889908, EPI_ISL_13889977, EPI_ISL_13890048, EPI_ISL_13890135, EPI_ISL_13890204, EPI_ISL_13890273, EPI_ISL_13890338, EPI_ISL_13890408, EPI_ISL_13890464, EPI_ISL_13890465, EPI_ISL_13890466, EPI_ISL_13890467, EPI_ISL_13890468, EPI_ISL_13890469, EPI_ISL_13890470, EPI_ISL_13890471, EPI_ISL_13890472, EPI_ISL_13890473, EPI_ISL_13890474, EPI_ISL_13890475, EPI_ISL_13890476, EPI_ISL_13890477, EPI_ISL_13890478, EPI_ISL_13890479, EPI_ISL_13890480, EPI_ISL_13890481, EPI_ISL_13890482 | | | |
| see above | Charité Universitätsmedizin Berlin, Institut für Virologie/Labor Berlin | Charité Universitätsmedizin Berlin, Institut für Virologie | Terry C. Jones, Julia Schneider, Barbara Mühlemann, Talitha Veith, Jörn Beheim-Schwarzbach, Julia Tesch, Marie Luisa Schmidt, Felix Walper, Tobias Bleicker, Caroline Isner, Frieder Pfäfflin, Ricardo Niklas Werner, Victor M. Corman, Christian Drosten |
| EPI_ISL_13891126 | Ministry of Health Turkey | Ministry of Health Turkey | Fatma Bayrakdar, Suleyman Yalcin, Gulay Korukluoglu |
| EPI_ISL_13908328 | Center of Diagnostics and Vaccine Development, Centers for Disease Control | Center of Diagnostics and Vaccine Development, Centers for Disease Control | Lin,J.-H., Chiu,S.-C., Huang,H.-I., Huang,W.-L., Fann,W.-B., Hsieh,P.-Y., Hsu,S.-C., Liu,P.-C., Chang,T.-Y. and Yang,J.-Y. |
| EPI_ISL_13908329, EPI_ISL_13908330, EPI_ISL_13908331, EPI_ISL_13908332, EPI_ISL_13908333, EPI_ISL_13908334, EPI_ISL_13908335, EPI_ISL_13908336, EPI_ISL_13908337, EPI_ISL_13908338, EPI_ISL_13908339, EPI_ISL_13908340, EPI_ISL_13908341, EPI_ISL_13908342, EPI_ISL_13908343, EPI_ISL_13908344, EPI_ISL_13908345 | | | |
| see above | Public Health Agency of Canada, National Microbiology Laboratory | Public Health Agency of Canada, National Microbiology Laboratory | Duggan,A., Hole,D., Yadav,C., Knox,N., Chapel,M., Tyler,A., Haidl,E., Domselaar,G.V., Antonation,K., Audet,J., Fernando,L., Hagan,M., Safronetz,D., Graham,M., Peters,G., Go,A., Laminman,V., Kaplen,B., Leung,A., Jolly,G., Fafard,J., Charest,H. and Levade,I. |
| EPI_ISL_13908346, EPI_ISL_13908347, EPI_ISL_13908348, EPI_ISL_13908349 | Centre for Biological Threats, Highly Pathogenic Viruses, Robert Koch Institute | Centre for Biological Threats, Highly Pathogenic Viruses, Robert Koch Institute | Brinkmann,A., Kohl,C., Pape,K., Uddin,S., Schrick,L., Michel,J., Jessen,H., Schaade,L. and Nitsche,A. |
| EPI_ISL_13908350, EPI_ISL_13908351, EPI_ISL_13908352, EPI_ISL_13908353, EPI_ISL_13908354, EPI_ISL_13908355, EPI_ISL_13908356, EPI_ISL_13908357, EPI_ISL_13908358, EPI_ISL_13908359, EPI_ISL_13908360, EPI_ISL_13908361, EPI_ISL_13908362, EPI_ISL_13908363, EPI_ISL_13908364, EPI_ISL_13908365 | | | |
| see above | Centre for Biological Threats, Highly Pathogenic Viruses, Robert Koch Institute | Centre for Biological Threats, Highly Pathogenic Viruses, Robert Koch Institute | Brinkmann,A., Kohl,C., Pape,K., Uddin,S., Schrick,L., Michel,J., Schaade,L. and Nitsche,A. |
| EPI_ISL_13953610, EPI_ISL_13953611 | Indian Council of Medical Research-National Institute of Virology | Indian Council of Medical Research-National Institute of Virology | Pragya Yadav, Rima Sahay, Anita Aich Shete, Sreelekshmy Mohandas, Priya Abraham |
| EPI_ISL_13955501 | Public Health Authority of the Slovak Republic | Laboratory of Genomics and Bioinformatics, Comenius University Science Park | Tomáš Szemes, Edita Staroová, Elena Tichá, Lucia Ševíková, Terézia Vrabová, Tatiana Sedláková, Miroslav Böhmer, Jaroslav Budiš, Pavol Mišenko |
| EPI_ISL_13958697 | Research and Evaluation, UKHSA | Research and Evaluation, UKHSA | Groves,N., Osman,K.L., Lewandowski,K.S., Carter,D.P., Pullan,S.T., Myers,R., Vipond,R. and Chand,M. |

| | | | |
|---|---|---|---|
| EPI_ISL_13958698 | Research and Evaluation, UKHSA | Research and Evaluation, UKHSA | Groves,N., Osman,K.L., Lewandowski,K.S., Pullan,S.T., Myers,R., Vipond,R. and Chand,M. |
| EPI_ISL_13983354, EPI_ISL_13983355 | Instituto de Infectologia Emilio Ribas | Instituto Adolfo Lutz Strategic Laboratory | Claudio Tavares Sacchi, Karoline Rodrigues Campos, Ariadne Ferreira Amarante, Marlon Benedito Nascimento Santos, Alex Domingos Reis, Adriano Abbud, Adriana Bugno |
| EPI_ISL_13983356 | INSPI-Centro de Referencia Nacional de Virus Exantemáticos, Gastroentéricos y Transmitido por Vectores. | INSPI-Dirección Técnica de Investigación, Desarrollo e Innovación INSPI-Centro de Referencia Nacional de Genómica, Secuenciación y Bioinformática | Andrés Carrazco-Montalvo, Diana Gutiérrez, Naomi Mora, Silvia Salgado-Cisneros, Johana Parrales-Valdiviezo, Martha Sánchez-Domenech, Diego Morales, Gulnara Borja-Cabrera, Leandro Patiño*. |
| EPI_ISL_13983888 | Bangkok Hospital Phuket | National Institute of Health, Department of Medical Sciences, Ministry of Public Health, Thailand | Pilailuk Okada; Siripaporn Phuygun; Nuttida Thongpramul; Thanutsapa Thanadachakul; Kazuhisa Okada; Archawin Rojanawiwat; Chakkarat Pitayawonganon; Supakit Sirilak |
| EPI_ISL_13993734, EPI_ISL_13993735, EPI_ISL_13993737, EPI_ISL_13993738, EPI_ISL_13993739 | California Department of Public Health | California Department of Public Health | Viral and Rickettsial Disease Laboratory |
| EPI_ISL_14003930 | University of Rochester Medical Center | University of Rochester Medical Center | Andrew Cameron, Mondraya Howard, Sara Connelly, Dwight Hardy, Kelly DeLary |
| EPI_ISL_14011193 | Bangkok Hospital Phuket | Thai Red Cross Emerging Infectious Diseases Clinical Center and Faculty of Medicine, Chulalongkorn University | Kusak Kukiattikoon, Waritta Dararattanaroj, Rome Buathong, Supaporn Wacharapluesadee, Sininat Petcharat, Ananporn Supataragul, Stefan Fernandez, Achawin Rojanaviwat, Chonticha Klungthong, Pilailuk Okada, Khajohn Joonlasak, Chakkarat Pitayawonganon, Opass Putcharoen |
| EPI_ISL_14021725 | Hosp. Municipal Enf. Antonio Policarpo de Oliveira | Instituto Adolfo Lutz Strategic Laboratory | Claudio Tavares Sacchi, Karoline Rodrigues Campos, Ariadne Ferreira Amarante, Marlon Benedito Nascimento Santos, Alex Domingos Reis, Adriano Abbud, Adriana Bugno |
| EPI_ISL_14033204, EPI_ISL_14033205, EPI_ISL_14033206, EPI_ISL_14033207, EPI_ISL_14033208, EPI_ISL_14033209, EPI_ISL_14033210, EPI_ISL_14033211, EPI_ISL_14033212, EPI_ISL_14033213 | Laboratory Medicine, UW Virology | Laboratory Medicine, UW Virology | Sereewit,J., Xie,H., Pavitra,R. and Greninger,A. |
| EPI_ISL_14049244, EPI_ISL_14049245 | Indian Council of Medical Research-National Institute of Virology | Indian Council of Medical Research-National Institute of Virology | Pragya Yadav, Rima Sahay, Anita Aich Shete, Sreelekshmy Mohandas, Priya Abraham |
| EPI_ISL_14050451, EPI_ISL_14050452, EPI_ISL_14050453, EPI_ISL_14050454, EPI_ISL_14050455, EPI_ISL_14050456, EPI_ISL_14050457, EPI_ISL_14050458 | Public Health Agency of Canada, National Microbiology Laboratory | Public Health Agency of Canada, National Microbiology Laboratory | Duggan,A., Hole,D., Yadav,C., Knox,N., Tyler,A., Haidl,E., Chapel,M., Domselaar,G.V., Graham,M., Audet,J., Fernando,L., Hagan,M., Safronetz,D., Leung,A., Peters,G., Go,A., Laminman,V., Kaplen,B., Antonation,K., Jolly,G., Griffiths,E., Charest,H., Levade,I. and Fafard,J. |
